# Behavioral dynamics of different stages of sexual motivation in male and female rats

**DOI:** 10.64898/2026.02.17.706347

**Authors:** John C. Oyem, Patty T. Huijgens, Julissa Mendoza, Roy Heijkoop, Eelke MS Snoeren

## Abstract

Sexual motivation is a complex concept involving both the initial drive to begin mating and the motivation to sustain copulation. Disruptions in sexual motivation are often observed in psychiatric disorders. This study proposes that sexual motivation can be divided into two distinct components: sexual incentive motivation and the drive to sustain copulation. To investigate this, we utilized the Motivation to Continue Copulation (MCC) test, which measures effort (nose pokes) to gain access to a sexual reward, and compared it with the Sexual Incentive Motivation (SIM) test and standard copulation tests. Male and female rats were trained on a fixed ratio (FR) 1 schedule using cheese rewards before transitioning to sexual rewards. After six FR1 sessions, the effort required increased to FR5 and progressive ratio (PR) schedules. Results revealed that sexual incentive motivation, measured by the SIM test, was higher in males after sexual experience, while females maintained consistent levels. In the MCC test, both males and females exhibited increased motivation to continue copulation with experience, but the motivation declined in the 2^nd^ ejaculatory series. These findings demonstrate that sexual motivation comprises distinct components. The MCC test effectively measures the drive to sustain copulation, while the SIM test assesses incentive motivation. This distinction is crucial for advancing behavioral neuroscience and understanding sexual dysfunction in psychiatric conditions.

## 1. Introduction

Motivation drives goal-directed behaviors and plays a crucial role in various psychiatric disorders. Disruptions in motivational processes are commonly observed in conditions such as depression, anxiety, schizophrenia, and addiction (Association, 2013; Everitt et al., 2008; Stone et al., 2008). When goal-directed behaviors involve a sexual stimulus, the motivation is specifically referred to as sexual motivation. Understanding sexual motivation is crucial, as it plays a central role in reproduction and is frequently disrupted in psychiatric and neurological conditions. In such disorders, sexual dysfunction can arise either from the underlying condition or as a side effect of its pharmacological treatment (Sewalem et al., 2022; Waldinger, 2015).

Over the past decades, the concept of sexual motivation has been extensively studied and revised, with many models distinguishing sexual arousal from sexual motivation. Sexual arousal depends on the activation of neural circuits controlling autonomic responses during sexual interaction (e.g., penile erection), whereas sexual motivation drives and maintains the various stages of sexual behavior (Bialy et al., 2019; Everitt, 1990; Le Moene and Agmo, 2019; Ventura-Aquino and Paredes, 2017; Ågmo, 2008; Ågmo and Laan, 2023, 2022). In this study, sexual motivation is proposed to consist of two distinct components: *sexual incentive motivation* and the *motivation to continue copulation*. Sexual incentive motivation represents the central motive state triggered by an incentive stimulus, increasing the likelihood of approach behavior. In contrast, the motivation to continue copulation reflects the drive to sustain copulatory activity, which is dynamically updated by sexual rewards during copulation. Although both components are influenced by e.g. physiological, hormonal, and cognitive factors, they operate during different phases of sexual interaction. Sexual incentive motivation is primarily influenced by the presence and value of sexual incentives before approach behavior, while the motivation to continue copulation emerges from the interplay between sensory feedback and the reward perception of copulatory interactions. Dissecting these components is crucial for understanding the complexity of sexual behavior and its underlying mechanisms, particularly in the context of psychiatric research where disruptions in motivational processes are common.

Sexual behavior in rats, both in the wild and in laboratory settings, is characterized by distinct patterns that differ between males and females. Male rats copulate in structured bouts consisting of repetitive sequences of mounts, intromissions, and ejaculations, while females display bouts of paracopulatory behaviors such as hops and darts, and respond to copulatory stimulation with lordosis (Beach, 1976, 1966; Dewsbury, 1967; Erskine, 1989). These primary parameters, along with derivative measures, have traditionally been used to quantify sexual motivation in rats (reviewed in (Heijkoop et al., 2018a)). For instance, higher numbers of paracopulatory behaviors are often interpreted as higher levels of sexual motivation (Beach, 1976; Gelez et al., 2013; Oyem et al., 2025; Snoeren et al., 2011a; Snoeren et al., 2011b), and shorter mount or ejaculation latencies are similarly considered indicators of greater sexual motivation (Everitt, 1990; Hull et al., 1990; Pfaus et al., 1990; Vega Matuszcyk et al., 1998). However, the complexity of motivation raises questions about what these parameters truly measure. As Barry J. Everitt rightly noted, ‘*a male with no legs will have endlessly long mount and intromission latencies, but in no way can be said to be deficient in sexual motivation or arousal*.’ ((Everitt, 1990), page 218). Furthermore, studies have shown that copulatory patterns and sexual incentive motivation can be manipulated independently of one another (Ellingsen and Ågmo, 2004; Heijkoop et al., 2018b; Huijgens et al., 2021b; Huijgens et al., 2024; Kippin et al., 2004; Le Moene and Agmo, 2019), making it difficult to interpret these traditional parameters in the context of specific motivational states. These observations underscore the need for a more detailed understanding of sexual behavior and the development of new tools to distinguish between different elements of sexual motivation.

Neuroscientists have developed several behavioral paradigms to study sexual motivation, with the Sexual Incentive Motivation (SIM) test being one of the most widely used. The SIM test measures time spent near an incentive stimulus versus a control stimulus, providing a straightforward assessment of sexual incentive motivation (Heijkoop et al., 2018b; Huijgens et al., 2023; Ågmo, 2003). This test is suitable for both sexually naïve and experienced animals, as it captures the central motive state before copulation is initiated (Portillo and Paredes, 2004; Snoeren et al., 2012; Ågmo, 2003). For example, sexually naïve male rats demonstrate immediate sexual incentive motivation for a receptive female or her odor, but sexual experience is required to recognize and respond to mixed or hidden odors (Ågmo, 2003). While the SIM test is effective for studying incentive-driven motivation, it does not address the motivation to sustain copulatory activity, which requires a different experimental approach.

To study the motivation to continue copulation, an operant response paradigm could be particularly suitable. In these tasks, animals are usually trained to perform specific actions, such as nose pokes or lever presses for a secondary reinforcer, like a light cue that was previously associated with sexual behavior (Everitt, 1990; Everitt and Stacey, 1987), or to gain access to a sexual partner (Bermant, 1961; French et al., 1972). This approach allows for the measurement of effort exerted to sustain sexual interactions. Fixed ratio (FR) schedules require animals to perform a set number of responses to gain access (Cummings and Becker, 2012; Matthews et al., 1997), while progressive ratio (PR) schedules gradually increase the response demands until the animal ceases to respond, providing a measure of the maximum effort an animal is willing to exert for a reward (Hodos, 1961; Richardson and Roberts, 1996). These paradigms have been used previously in motivational research and could be employed for studying motivation to continue copulation as well.

This study had three primary aims: (1) to establish whether the operant response task is effective for studying the motivation to continue copulation, (2) to characterize the differences in sexual motivation between sexually naïve and experienced rats, and between male and female rats, and (3) to evaluate whether parameters from the standard copulation (COP) test could serve as measures of sexual incentive motivation or the motivation to continue copulation. By employing the SIM and motivation to continue copulation (MCC) tests, this study provides a framework for dissecting the distinct components of sexual motivation.

## 2. Materials and Methods

### 2.1 Animals

The study was conducted using sexually naïve male (N = 34) and female Wistar rats (N = 34), each weighing approximately 200g. These experimental animals were obtained from the Janvier laboratory in France. Rats were housed in same-sex pairs in Macrolon IV ® homecages under a reversed 12 h light/dark cycle (lights on between 23:00 and 11:00) at 21 ± 1 °C temperature and controlled humidity of 55 ± 10 %. The rats had unrestricted access to standard rodent food pellets (low phytoestrogen maintenance diet, #V1554, Ssniff, Germany) and water *ad libitum*. All procedures in this experiment complied with European Union Council Directive 2010/63/EU and were performed in accordance with the Norwegian Food Safety Authority with the ethical approval number FOTS ID 30239 approved on the 1^st^ of June 2023.

### 2.2 Drugs

Silastic capsules (medical grade Silastic tubing, length 5mm, 0.0625 inch inner diameter, 0.125 inch outer diameter, Degania Silicone, Degania Bet, Israel) containing 10 % 17β-estradiol (Sigma, St. Louis, USA, product Nr: E-8515) in cholesterol (Sigma, St. Louis, USA, product Nr: C3292) were made for the female Wistar rats. The two ends of the silastic tubing were closed off by inserting pieces of toothpick and sealed off with medical-grade adhesive silicone (NuSil Silicone Technology, Carpinteria, USA). Progesterone solution was made by dissolving progesterone powder (Sigma, St. Louis, USA, product Nr: P-0130) in peanut oil (Apotekproduksjon, Tromsø, Norway) at a 2.5 mg/mL concentration, from which 0.2 mL was administered subcutaneously 4 hours before testing (Oyem et al., 2025; Snoeren et al., 2011a).

### 2.3 Ovariectomy

After one week of acclimatization to the animal facility, all female rats were ovariectomized following the standard methods (Ågmo, 1997). Rats were anesthetized with isoflurane and placed on a sterile operating table. The skin was shaved and disinfected, after which analgesia was administered subcutaneously (0.05 mg/kg Temgesic & 2 mg/kg Metacam). A mediodorsal incision of about 1 cm in length was made on the skin, followed by a smaller incision of about 0.5 cm in the muscular layer to expose the peritoneal layer. Next, the ovaries were identified and surgically removed, and the fallopian tubes were ligated and gently placed back in the posterior abdomen. The incised muscle layer was closed with absorbable sutures (Vicryl 4–0, Ethicon). A silastic capsule containing 10 % 17β-estradiol was inserted subcutaneously, and finally, the skin was closed using wound clips. For postoperative care, all females were administered analgesic treatment, which consisted of 2 mg/kg Metacam and 0.05 mg/kg Temgesic in 24 and 48 hours while recuperation was closely monitored. All ovariectomized female rats were given a recovery period of two weeks before the start of behavioral testing.

### 2.4 Experimental design

Before the start of the experiments, the stimulus male (N = 14) and female (N = 14) rats were sexually trained in three copulation tests for 30 minutes. The experimental male (N=20) and female (N=20) rats were trained in the motivation to continue copulation set-up with cheese rewards until they successfully learned to associate nose poking with door opening, consequently allowing access to the reward (see Fig. 1A and motivation to continue copulation test for more details).

**Figure 1.**
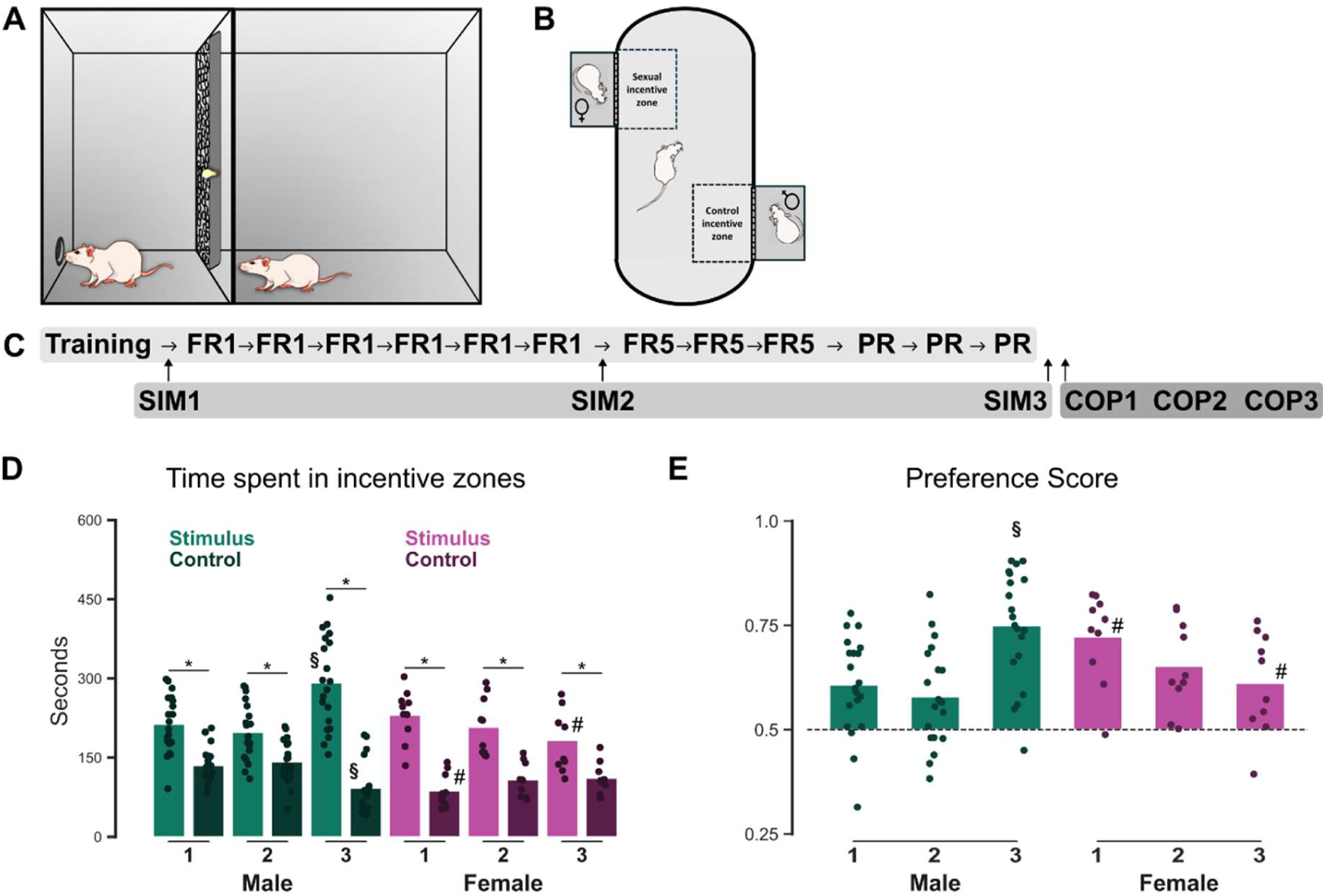
A) Schematic illustration of the Motivation to Continue Copulation (MCC) test box, B) Schematic illustration of the Sexual Incentive Motivation (SIM) test with male subject as example, C) Schematic overview of the timeline of the experiment. SIM1-3 = sexual motivation test 1 to 3, FR1 = MCC test on fixed ratio 1, FR5 = MCC on fixed ratio 5, PR = MCC on progressive ratio, COP1-3 = copulation test 1 to 3. D) Time spent in the sexual incentive (stimulus) versus control zone measures in the 1^st^, 2^nd^ and 3^rd^ SIM test, E) Preference score in the 1^st^, 2^nd^ and 3^rd^ SIM test (all significantly different from 0.05 (no preference)). Data is shown for both male (green) and female (purple) rats in dots as individual datapoints and bars for the mean. * p≤ 0.05 between the connected groups, § p≤ 0.05 compared to SIM1 and SIM2 of the same category, ^#^ p≤ 0.05 compared to male rats of the same category.

After successful training in the motivation to continue copulation test box, each experimental (and sexually naïve) rat underwent a series of behavioral test sessions in three different tests: the sexual incentive motivation test (SIM, Fig. 1B), the motivation to continue copulation test (MCC) and the copulation test (COP). The male rats were tested at a two-day interval, while the female rats were tested at a four-day interval to simulate the female rats’ natural estrous cycle (Snoeren, 2019).

As shown in Figure 1C, all experimental rats started with a SIM test (SIM 1), after which they were tested in a series of six MCC tests on a fixed ratio 1 (FR1) schedule to gain physical access to a sexually experienced partner of the opposite sex. Next, we conducted a second SIM test (SIM 2), which was followed by three MCC tests on a fixed ratio 5 (FR5) schedule and another three MCC tests on a progressive ratio (PR) schedule. Finally, all experimental (and sexually experienced) rats were tested in the last SIM test (SIM 3), followed by three standard copulation tests (COP 1 – COP 3). The copulation tests of the male rats were performed in a single-compartment copulation box, while the females underwent paced mating in a two-compartment copulation box (more details under behavioral testing)

All tests were conducted during the dark period of the day. The order of rats to be tested in each experiment was randomly counterbalanced across days. All behavioral tests were recorded with a digital video camera (Sony HDR-CX240E handycam), and the behavior was manually annotated afterwards using Observer 17XT software (Noldus, Wageningen, The Netherlands).

### 2.5 Behavioral testing

#### 2.5.1 Sexual incentive motivation test (SIM)

Used to study sexual incentive motivation (Huijgens et al., 2023; Huijgens et al., 2021b; Huijgens et al., 2024; Snoeren et al., 2012; Snoeren and Ågmo, 2014; Ågmo, 1997, 2003), the SIM apparatus consisted of a rectangular open field arena of 100 x 50 x 45 cm diameter with oval-shaped corners placed in a dimly lit (5 lx) room (Fig, 1B). On both long sides of the box two stimulus incentive boxes (25 x 10 x 25cm) were attached to the arena. For male subject rats, a receptive female was used as a sexual stimulus and a male rat as a (social) control stimulus. For the female subject rats, on the other hand, an intact male rat acted as a sexual stimulus, while a female was used as a control (not injected with progesterone).

The control and sexual stimuli were placed in each stimulus box for 5 minutes before the start of each test. A steel mesh separated these stimulus boxes from the arena, allowing interaction between the rats without physical contact. The subject rats were habituated to the arena for three days before the start of the test by placing them in the middle of the open arena (without stimuli being present) for 10 minutes.

At the start of the test, subject animals, either male or female rats, were placed in the middle of the arena and then video-tracked with a video camera located above the test boxes using Ethovision software (Noldus, Wageningen, the Netherlands) for 10 minutes. Two virtual incentive areas measuring 20 x 30 cm were mapped out in front of each stimulus box in the arena. The subject was considered to be in this area whenever its center of gravity is within the zone. We randomly interchanged stimulus boxes across tests to rule out any influence of spatial location. Several parameters were measured in the SIM test, including time spent in each of the incentive zones, mean velocity, total distance, and total duration moved. In addition, a preference score was calculated as the ratio of time spent in sexual incentive zones to the total time in both the sexual and control incentive zones. The SIM boxes were carefully cleaned after each experimental day.

#### 2.5.2 Motivation to continue copulation test (MCC)

The MCC test box consisted of two different compartments: the nose poke compartment (30×30×70 cm) and the reward compartment (45×30×70 cm, Fig. 1A). The two compartments were separated by a wall with two doors opening horizontally (one on the left and one on the right side of the wall) that could be operated both manually and automatically. In the manual control, both doors opened simultaneously, while in the automatic control only the right door opened. These doors were perforated with tiny holes that allow communication between the subject rat and the sexual stimulus via visual, auditory, and olfactory cues. A light positioned on the wall of the dividing doors facing the nose poke compartment signaled a successful nose poke trial and the opening of the door 5 seconds later. In the nose poke compartment, a nose poke device was located at the lower portion of the wall opposite to the right door. The nose poke device was equipped with a yellow light that was on when the device was active. The floor of both compartments was covered with wood chip bedding.

All subject rats were first habituated in the MCC test box for 10 minutes in two sessions (without a reward being present), followed by training sessions in which they learned to associate nose poking with the door opening for access to a food reward (small pieces of cheese). The training sessions were conducted daily for about 6 - 13 days until the animals had learned to poke for access consistently. The training consisted of three phases:

Phase 1: the subject rats were placed in the MCC test box (in the nose poke compartment) for 30 minutes, while the experimenter manually controlled the doors and reward delivery. Whenever the rat was in proximity to the nose poke device, the experimenter opened the door and dropped some pieces of cheese near the rat. The experimenter gradually dropped the cheese closer to the door until it was dropped in the reward compartment.

Phase 2: the subject rats were placed in the MCC test box for 30 minutes during which the box was manually controlled. During this phase, the rat had to nose poke to gain access to the cheese reward in the other compartment. Every time the rat nose poked, the experimenter manually opened the door and dropped a piece of cheese in the reward compartment. The subject rat spent 15 seconds eating before being placed back in the nose poke compartment for a new round.

Phase 3: In the final training phase, the box was automatically controlled and the rats had to perform the task themselves on a fixed ratio 1 schedule. Upon every nose poke, the light turned on, and 5 seconds later the left door opened. The door remained open for 15 seconds, allowing the rat to cross and eat a piece of cheese in the other compartment. After the door automatically closed again, the rat was manually placed back in the nose poke compartment and a new round of pokes to gain access to the cheese reward was started.

Once all rats had successfully learned the operant response, the experimental phase was started. The procedures remained the same as phase 3 of the training, except that a sexual reward replaced the cheese reward. It should be mentioned that both the subject and stimulus rats were now able to cross to the other compartment when the door opened. For the test, it does not matter in which compartment they had their physical interaction.

To begin the (experimental) MCC test, the subject rats were placed in the nose poke compartment while the stimulus rat of the opposite sex (sexual reward) was placed in the reward compartment. At this point, the sliding doors were closed. After 3 minutes of no physical contact interaction between the subject and the stimulus rat, the light in the nose poke device turned on signaling an active nose poke device. Whenever a rat made a successful nose poke, the light above the door turned on for 5 seconds, followed by the opening of the left door. The left door stayed open for 15 seconds to allow the subject rat to physically interact with the sexual reward. During this period the light in the nose poke device was turned off, and no further nose pokes were recorded. Fifteen seconds after physical contact possibility, the subject and stimulus rats were placed back into their respective compartment until the subject rat made another successful nose poke to gain new access for physical interaction. For the female subject rats, the male stimulus rat was replaced by another male after an ejaculation was achieved. The female stimulus for the male subject rats remained the same for the full duration of the test.

The subject rats underwent multiple tests with three different nose poke schedules to gain access to the sexual reward. They started with six FR1 tests, followed by three FR5, and 3 PR tests.

- Fixed ratio 1 (FR1): one nose poke to open the door.
- Fixed ratio 5 (FR5): five nose pokes to open the door.
- Progressive ration (PR): The number of nose pokes needed to open the door increased progressively (1, 2, 4, 6, 9, 12, 15, 20, 25, etc. as used by (Richardson and Roberts, 1996))

All FR1 and FR5 tests were terminated after 30 minutes had elapsed from the start of the test session or when two ejaculations were obtained. All PR tests were terminated following 10 mins of inactivity (no nose pokes) or 30 minutes had elapsed from the start of the test session. The exception to this protocol was the third PR test, which was terminated after one hour of testing (keeping the other conditions the same).

The successful nose pokes were registered by an Arduino system that automatically controlled the lights and door of the MCC test box. The sessions were recorded on video and afterwards manually scored by a trained observer using the Observer 17XT software (Noldus, Wageningen, The Netherlands).

#### 2.5.3 Copulation test (COP)

##### 2.5.3.1 Single-compartment copulation test (males)

The copulation test of the male subject rats was performed in a single-compartment copulation test box. This box consisted of a rectangular steel box measuring 40 x 60 x 40 cm with a Plexiglas front and was conducted in a room with dim lights (ca 5 lux). In this test, a male subject rat was placed in the copulation box for 5 mins after which a (hormonally primed, sexually trained) receptive female was introduced. The test sessions lasted for 30 minutes. The sessions were recorded on video and afterwards manually scored by a trained observer using the Observer 17XT software (Noldus, Wageningen, The Netherlands).

##### 2.5.3.1 Two-compartment copulation test (females)

The copulation test of the female subject rats was performed in a two-compartment copulation test box, allowing the females to pace their sexual interactions. The apparatus is the same as the single-compartment box, except that the interior space is divided into two compartments by a transparent compartment divider. The divider has three 4 cm diameter holes at the bottom, which serve as an escape route for the females during copulation. As a result, one compartment (40 x 45 x 40 cm) is available for both the male and female, while a smaller compartment (40 x 15 x 40 cm) can only be accessed by the female.

In this test, a receptive female rat was placed into the female compartment of the copulation apparatus and was allowed to habituate for 5 minutes. During this period, the female was able to cross to the other compartment. After habituation, a sexually experienced male rat was introduced in the male compartment. They were allowed to copulate for 30 minutes. Every time a male rat achieved an ejaculation, an experimenter replaced the stimulus male with another sexually experienced male rat. The sessions were recorded on video and afterward manually scored by a trained observer using the Observer 17XT software (Noldus, Wageningen, The Netherlands).

### 2.6 Behavioral analysis

For each test, different kinds of behaviors were annotated, and parameters were calculated (see Table 1). For the MCC test, we manually scored the correct nose pokes, door openings, 1^st^ crossing after door opening of either subject or stimulus rat, number of paracopulatory behaviors (females only), lordosis responses (females only, scored on a scale of 0-3 with 0 being no lordosis response (Hardy and Debold, 1971)), mounts, intromissions, and ejaculations. In addition, in order to calculate nose poke bouts, we also scored attention away from the nose poke device. A nose poke bout was determined as a cluster of nose pokes within a set of nose pokes required for a successful door opening with continuous attention to the nose poke device. Attention away from the nose poke device interrupts the nose poke bout and starts a nose poke time-out. As soon as a new nose poke was performed, a new nose poke bout was started. From these behavioral annotations, we also calculated additional parameters, such as nose poke intervals, in three different ways (see Table 1). In the PR tests, we also calculated the breakpoint, which was set as the last completed set of nose pokes required for a successful door opening after 10 minutes without nose pokes or the end of the test.

**Table 1.** Behavioral parameters description.

| Behavioral outcome | Measure |
| --- | --- |
| Number of nose pokes (MCC) | Total number of nose pokes during the test. |
| Nose poke latency (MCC) | Latency to the first nose poke after returning to the nose poke chamber. |
| Contact latency (MCC) | Latency to make contact with stimulus after a successful door opening by crossing to the other chamber. |
| Number of crossings (MCC) | Number of times contact with stimulus was initiated after door opening by crossing to the other chamber. |
| Nose poke interval (MCC) | Time from 1<sup>st</sup> to last nose poke needed to open the door divided by the number of nose pokes. |
| Breakpoint (MCC) | Last completed set of nose pokes required for a successful door opening after 10 minutes without nose pokes. |
| Nose poke bout (MCC) | A cluster of nose pokes within a set of nose pokes required for a successful door opening with continuous attention for the nose poke device. Attention away from the nose poke device interrupts the nose poke bout. |
| Nose poke time-out (MCC) | A break within a set of nose pokes starting when attention was drawn away from the nose poke device and ending with the next nose poke. |
| Latency to the first paracopulatory behavior, mount, or intromission (MCC, COP) | Latency to the first paracopulatory behavior, mount, intromission, or ejaculation from the start of the test. |
| Number of paracopulatory behaviors, mounts, intromissions, and ejaculations (MCC, COP) | Total number of paracopulatory behaviors, mounts, intromissions, and ejaculations (performed or received). |
| Number of copulation (MCC, COP) | Sum of mounts, intromissions, and ejaculations (performed or received). |
| Ejaculation latency (MCC, COP) | Latency to ejaculation from the first paracopulatory behavior, mount or intromission (performed or received). |
| Lordosis quotient (MCC, COP) | Number of lordosis responses divided by the number of received copulations multiplied by 100%. |
| Lordosis score (MCC, COP) | Lordosis quality measured on a scale of 0-3 (0 is no lordosis). The lordosis score is calculated by the sum of lordosis scores divided by the number of received copulations. (Hardy and Debold, 1971) |
| Intromission ratio (COP) | Number of intromissions divided by the total number of copulatory behaviors (mounts + intromissions) |
| Inter-intromission interval (COP) | Ejaculation latency divided by number of intromissions |
| Post-ejaculatory interval (COP) | Time from an ejaculation to the next copulatory behavior (mount or intromission) |
| Percentage of exits after mount, intromission, and ejaculations (COP) | Percentage of withdrawals to the female compartment within 5 seconds after receiving a mount, intromission, or ejaculation. |
| Contact return latency after mount, intromission, and ejaculation (COP) | The mean duration for the females to re-enter the male compartment after she withdraws upon a mount, intromission, or ejaculation. |
| Number of sexual bouts (COP) | The total number of sexual bouts defined as a series of behaviors that begins with either a paracopulatory behavior or lordosis response (females) or a mount, intromission, or ejaculation (males) and continues until a behavior was displayed that was not oriented towards the sexual partner. The last paracopulatory behavior or lordosis response (females) or a mount, intromission, or ejaculation (males) marked the end of a sexual bout, and the start of a time-out consisting of behaviors that are not oriented at the sexual partner. (Huijgens et al., 2021a; Oyem et al., 2025) |
| Sexual bout duration (COP) | Time from the first sexual behavior in a sexual bout until the last behavior within the sexual bout (Huijgens et al., 2021a; Oyem et al., 2025) |
| Time-out duration (COP) | Time from the end of one sexual bout to the start of the next sexual bout. (Huijgens et al., 2021a; Oyem et al., 2025) |
| Time spent in zones (SIM) | Time spent by subject rats in each incentive zone. |
| Preference score (SIM) | Time spent in incentive stimulus zone divided by the total time spent in both the incentive stimulus and control zones. |

*Note: MCC refers to the motivation to continue copulation test, COP to the copulation test and SIM to the sexual incentive motivation test. Some parameters are used in multiple tests.*

For the copulation test, we annotated the same (para)copulatory behaviors as in the MCC test. From these measures, we also calculated the intromission ratio, inter-intromission interval, and post-ejaculatory interval (see Table 1, reviewed in (Heijkoop et al., 2018b)). For the females, we also recorded the escapes to the female compartment after mounts, intromissions and ejaculations within 5 seconds, so that we could calculate the percentages of exits and the contact return latencies. Furthermore, we registered the attention away from the sexual stimulus during copulation in order to calculate the sexual bout parameters. For the males, a sexual bout was set as a series of mounts, intromissions, and/or ejaculations that were uninterrupted by other behaviors that were not oriented towards the female. As soon as the attention turned away from the female, the last mount, intromission, or ejaculation signaled the end of a sexual bout, and the start of a time-out (Huijgens et al., 2021a). For the female, a sexual bout was calculated similarly, except that it now started and ended with either a paracopulatory behavior or a lordosis response (Oyem et al., 2025).

The location of the rat in the sexual incentive motivation test was automatically tracked. From these coordinates, we measured the time spent in each incentive zone and calculated the preference score as the time spent near the sexual stimulus divided by the time spent in both incentive zones. In addition, ambulatory activity was explored in the form of the total distance moved during the test, the mean velocity of movement while moving, and the time spent moving.

### 2.7 Statistical analysis

A customized Python script was developed to analyze all data and generate the figures. For parameters with multiple data points per animal (e.g., latency to poke, time-out duration, nose poke interval, etc.), the average value for each animal was calculated and used as its final data point for further group analysis. Statistical analyses were performed using SPSS software (version 29, IBM, Armonk, USA), with the threshold for statistical significance set at *p* ≤ 0.05. A Shapiro-Wilk test confirmed that the data were not normally distributed. However, due to the complexity of the study design and the suitability of the test for such data, a linear mixed model was employed. This model included the factors *Sex*, *Test*, and *Sex*Test* for the overall test analysis, and the factors *Sex*, *Test*, *Series*, and *Sex*Test*Series* for the analysis comparing ejaculatory series 1 and 2.

For the SIM test, the factors *Sex*, *Test*, *Incentive zone*, and *Sex*Test*Incentive zone* were used to compare the time spent in the incentive zone. In cases where a significant interaction effect was observed, a Bonferroni-corrected post hoc test was applied. Additionally, the preference score had to exceed 0.5 (indicating no preference, i.e., equal time spent in the incentive and no-incentive zones) for the incentive to be considered effective. A *t*-test was used for statistical evaluation in these cases. Finally, when analyzing the relationship between the duration of sexual bouts, time-outs, and other parameters, the Spearman correlation test was employed. To prevent false positive outcomes, a conservative approach was used in which correlations needed to appear in a majority of the test, and have p-values under 0.01.

## 3. Results

### 3.1 Sexual incentive motivation

During the 10 minute exposure to a sexual incentive versus social control stimulus, we found that both male and female subject rats spent significantly more time with the sexual incentive than with the control stimulus in all three SIM tests (Fig 1D, interaction effect Sex*Test*Incentive zone: F_(2,140)_=12.654, p<0.001, Female: SIM1: p<0.001, SIM2: p<0.001, SIM3: p=0.004, Males: SIM1: p<0.001, SIM2: p<0.001, SIM3: p<0.001). In addition, we found a significant preference score compared to 0.5 (no preference) in all SIM tests in both male and female rats (Fig. 1E, Female SIM1: *t*_(9)_=6.568, p<0.001, Female SIM2: *t*_(9)_=6.568, p<0.001, Female SIM3: *t*_(9)_=6.568, p<0.001, Male SIM1: *t*_(19)_=4.084, p<0.001, Male SIM2: *t*_(19)_=2.975, p=0.008, Male SIM3: *t*_(19)_=8.360, p<0.001). Interestingly, when sexually naïve (SIM1) versus experienced rats (SIM3) were compared, we found that only male rats significantly increased preference score when gaining experience (interaction effect Sex*Test: F_(2,56)_=10.186, p<0.001, SIM3 vs SIM1 and SIM2, p<0.001). Finally, a sex difference was found in the 1^st^ and 3^rd^ SIM tests with regard to the preference score. While females had a higher preference score for the incentive stimulus than males in SIM1 (p=0.015), the sexual incentive motivation in males was larger than in females in SIM3 (p=0.004, Fig. 1E).

### 3.2 Motivation to continue copulation

Cheese rewards were initially used to train both male and female rats in the MCC test box, where they learned to perform nose pokes on a FR1 schedule to gain access to the reward. Following successful training, the sexually naïve rats were transitioned to testing with a sexual reward, where a nose poke allowed access to a mate partner. The rats underwent six test sessions on the FR1 schedule, during which they gained sexual experience. After completing the six FR1 tests, the rats continued with three test sessions on a FR5 schedule, requiring increased effort to gain access to the sexual reward, followed by three additional test sessions on a PR schedule, where the effort (number of nose pokes) required to gain access to the mate progressively increased (Fig. 1C).

#### 3.2.1 FR1

As shown in Fig. 2A, no relevant differences in the number of nose pokes were observed across the six FR1 test sessions for either male or female rats. Additionally, there was no overall sex difference in the number of nose pokes (Sex (F_(1,38)_=1.155, p=0.289), Test (F_(5,190)_=3.127, p=0.010) and Sex*Test interaction effects (F_(5,190)_=1.445, p=0.210)). Only during the 4th FR1 test (FR1-4), male rats poked significantly more often than in the 1st FR1 test (FR1-1). Since this effect was not consistent across tests, it is likely an artifact rather than a meaningful result. In contrast, the latency to nose poke decreased over test sessions in male, but not in female rats (Fig. 2B). Statistical analysis revealed significant effects of Sex (F_(1,38.174)_=7.671, p=0.009) and Test (F_(5,186.972)_=8.688, p<0.001), but the Sex*Test interaction was not significant (F_(5,186.972)_=1.998, p=0.081)). For female rats, the latency to nose poke in FR1-1 was only significantly longer than in FR1-5 (p = 0.045). In male rats, however, the latency to nose poke decreased more consistently, with significant reductions observed after the 3rd FR1 test (FR1-1 vs FR1-4: p=0.001, FR1-1 vs FR1-5: p=0.037, FR1-1 vs FR1-6: p<0.001, FR1-2 vs FR1-3: p=0.014, FR1-2 vs FR1-4: p<0.001, FR1-2 vs FR1-5: p=0.007, FR1-2 vs FR1-6: p<0.001). Only in FR1-2, the latency to poke was significantly longer in males compared to females (p < 0.001). After a successful nose poke, the door opened and allowed for crossing to the reward chamber. Male and female rats crossed on nearly all occasions to gain access to the mate, and no significant differences were found between sexes or across tests (Fig. 1C, Sex (F_(1,38)_=2.664, p=0.111), Test (F_(5,190)_=0.451, p=0.812) and Sex*Test interaction effects (F_(5,190)_=1.326, p=0.255)). Similarly, no differences were observed in the latency to cross to the reward chamber (Fig. 1D, Sex (F_(1,35.359)_=0.004, p=0.953), Test (F_(5,176.410)_=0.538, p=0.748) and Sex*Test interaction effects (F_(5,176.410)_=1.580, p=0.168)). Altogether, these findings suggest that gaining sexual experience only shortens the latency to nose poke but does not influence sexual motivation to continue copulation or any other parameters under an FR1 schedule.

**Figure 2.**
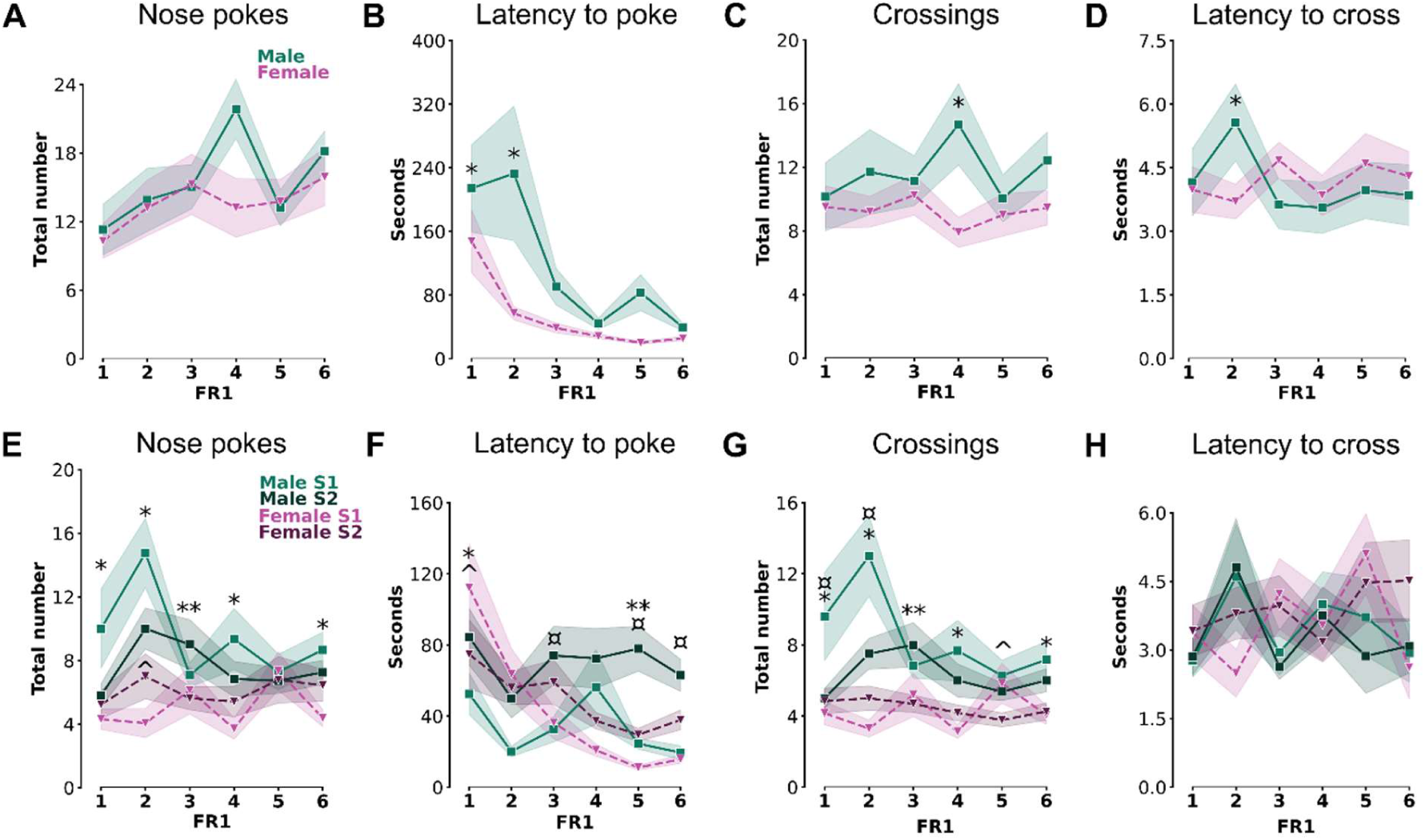
A) Total number of nose pokes, B) Latency to nose poke, C) Total number of crossings, and D) Latency to cross during the complete test under a FR1 schedule, and E) Total number of nose pokes, B) Latency to nose poke, C) Total number of crossings, and D) Latency to cross during the 1st (S1) and 2^nd^ (S2) ejaculatory series under a FR1 schedule. Data are shown of male (green line) and female (purple dotted line) rats in A-D, and male (green line) and female (purple dotted line) rats per ejaculatory Series 1 (light color line) and Series 2 (dark color dotted line) in E-H. Shadows represent the SEM. * p≤ 0.05 difference between male and female rats in total test (A-D) or Series 1 (E-H), ** p≤ 0.05 difference between male and female rats in Series 2, ^ p≤ 0.05 difference between Series 1 and 2 of female rats, ¤ p≤ 0.05 difference between Series 1 and 2 of male rats.

The next question is whether the motivation to continue copulation increases or declines after an ejaculation has been achieved. If this is the case, such an effect should be observable in the MCC test when comparing the 1st and 2nd ejaculatory series (S1 and S2, respectively) in rats who obtained two ejaculations. We found a significant Sex (F_(1,42.505)_=28.417, p<0.001) and Sex*Test*Series interaction effects (F_(16,246.975)_=1.872, p=0.024) on the number of nose pokes per series, but not an effect on Test (F_(5,259.887)_=1.525, p=0.182) or Series (F_(1,236.278)_=0.578, p=0.448). Post-hoc analysis, however, revealed that female rats only poked more during S2 than S1 in FR1-2 (Fig. 2E, p=0.037), while no differences in number of nose pokes were found in male rats in the six FR1 tests. During S1, however, male rats generally had a higher number of nose pokes than female rats, which was significantly different in FR1-1 (p=0.008), FR1-2 (p<0.001), FR1-4 (p=0.004), and FR1-6 (p=0.006). During S2, this was only found in FR1-3 (p=0.041). Despite the same number of nose pokes across series, there was an effect on the latency to poke (Fig. 2F, Sex (F_(1,41.573)_=0.656, p=0.423), Test (F_(5,257.317)_=7.535, p<0.001), Series (F_(1,236.382)_=19.217, p<0.001) and Sex*Test*Series interaction effects (F_(16,244.339)_=3.094, p<0.001). While female rats only had a longer latency in S1 compared to S2 in FR1-1 (p=0.009), male rats on the other hand, took longer time to poke in S2 compared to S1 in FR1-3 (p=0.008), FR1-5 (p<0.001) and FR1-6 (p=0.004). A sex difference in poke latency was found in FR1-1, with females having longer poke latencies in S1 than males (p=0.002), and FR1-5 with longer poke latencies in males than females during S2 (p=0.001). While the poke latencies clearly decreased over time in females (S1 FR1-1 vs FR1-2: p=0.003, FR1-1 vs FR1-3: p<0.001, FR1-1 vs FR1-4 p<0.001, FR1-1 vs FR1-5: p<0.001, FR1-1 vs FR1-6: p<0.001, FR1-2 vs FR1-4: p=0.006, FR1-2 vs FR1-5: p<0.001, FR1-2 vs FR1-6: p=0.004; S2 FR1-1 vs FR1-4: p=0.037m FR1-1 vs FR1-5: p=0.005), no such effect was found in males. Notably, only rats that achieved two ejaculations were included in the between-series analysis.

Consequently, the sluggish rats that may have contributed to the prolonged total latencies observed in FR1-1 and FR1-2 of Figure 2B might be excluded from FR1-1 and FR1-2 of Figure 2F, potentially explaining the absence of latency declines in males. Similarly, it should be noted that the post-ejaculatory interval is included in S2 which theoretically could also influence motivation and thus latency to poke.

While the number of nose pokes was not significantly different between S1 and S2, the males did cross more often in S1 than S2 in FR1-1 and FR1-2 (Fig. 2G, Sex (F_(1,36.158)_=37.996, p<0.001), Test (F_(5,258.795)_=2.070, p=0.070), Series (F_(1,232.881)_=7.006, p=0.009) and Sex*Test*Series interaction effects (F_(16,243.652)_=2.652, p<0.001), Post-hoc S1 vs S2: Males: FR1-1: p=0.009, FR1-2). Female rats, on the other hand, did not cross more or less frequently, except for a decrease in crossings in S2 compared to S1 in FR1-5 (p=0.026). In contrast, no differences were found in the latency to these crossings between S1 and S2 or male and female rats (Fig. 2H, Sex (F_(1,29.871)_=0.710, p=0.406), Test (F_(5,254.876)_=0.797, p=0.553), Series (F_(1,223.804)_=0.068, p=0.795) and Sex*Test*Series interaction effects (F_(16,236.408)_=1.138, p=0.320)).

#### 3.2.1 FR5

After the completion of six FR1 tests, the effort to gain access to the sexual stimulus was increased. In the next three tests, rats had to poke five times to open the door (FR5). Under this regime, no differences were found in the number of nose pokes (Fig. 3A, Sex (F_(1,38)_=0.931, p=0.341), Test (F_(2,76)_=3.637, p=0.031) and Sex*Test interaction effects (F_(2,76)_=0.551, p=0.578)), or number of crossings (Fig. 3C, Sex (F_(1,38)_=2.940, p=0.095), Test (F_(2,76)_=1.242, p=0.295) and Sex*Test interaction effects (F_(2,76)_=0.363, p=0.697)). Similarly, there was no difference in latency to nose poke (Fig. 3B, Sex (F_(1,38.134)_=14.478, p<0.001), Test (F_(2,75.364)_=0.047, p=0.954) and Sex*Test interaction effects (F_(2,75.364)_=1.314, p=0.275) or cross (Fig. 3D, Sex (F_(1,35.173)_=5.817, p=0.021), Test (F_(2,69.650)_=0.879, p=0.420) and Sex*Test interaction effects (F_(2,69.650)_=0.596, p=0.554) between the test. However, male rats did have a significantly longer latency to nose poke than females in all FR5 tests (FR5-1: p=0.021, FR5-2: p=0.001, FR5-3: p<0.001), and a shorter latency to cross in the FR5-2 test (p=0.020).

**Figure 3.**
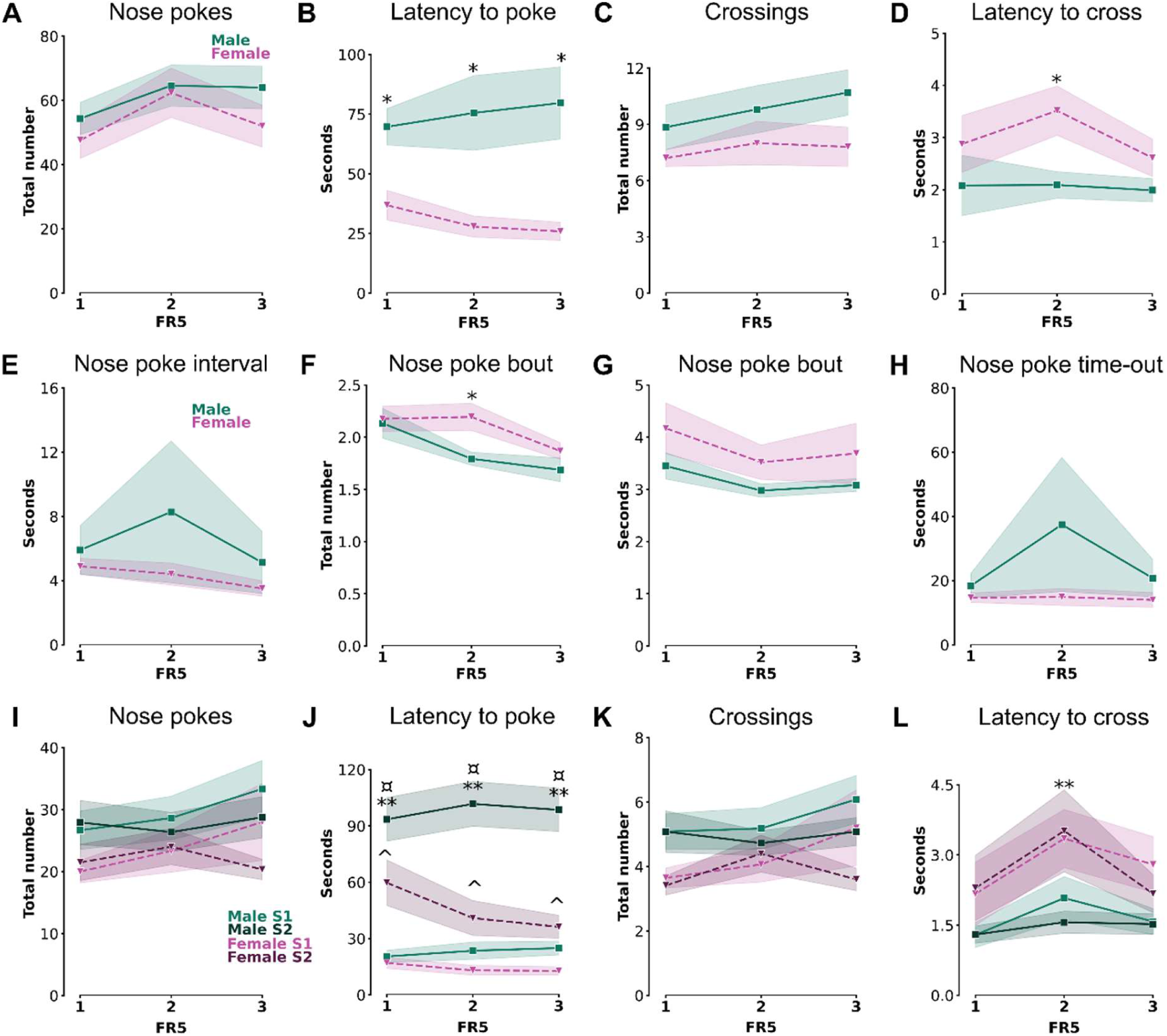
A) Total number of nose pokes, B) Latency to nose poke, C) Total number of crossings, and D) Latency to cross, E) Nose poke interval, F) Total number of nose poke bouts, G) Mean duration of nose poke bouts, and H) Mean duration of nose poke time-outs during the complete test under a FR5 schedule, and I) Total number of nose pokes, J) Latency to nose poke, K) Total number of crossings, and L) Latency to cross during the 1st (S1) and 2nd (S2) ejaculatory series under a FR5 schedule. Data are shown of male (green line) and female (purple dotted line) rats in A-D, and male (green line) and female (purple dotted line) rats per ejaculatory Series 1 (light color line) and Series 2 (dark color dotted line) in E-H. Shadows represent the SEM. * p≤ 0.05 difference between male and female rats in total test (A-D) or Series 1 (E-H), ** p≤ 0.05 difference between male and female rats in Series 2, ^ p≤ 0.05 difference between Series 1 and 2 of female rats, ¤ p≤ 0.05 difference between Series 1 and 2 of male rats.

To study the nose poke behavior in more detail, we calculated the nose poke intervals and divided the nose pokes into nose poke bouts. A nose poke bout consisted of a sequence of nose pokes that were performed without interruptions in attention. As soon as the rat directed its attention away from the nose poke device, a time-out began and lasted until a new nose poke bout was started. The data revealed that there was no difference in nose poke intervals between tests and sex (Fig. 3E, Sex (F_(1,38.136)_=0.875, p=0.355), Test (F_(2,75.404)_=0.803, p=0.452) and Sex*Test interaction effects (F_(2,75.404)_=0.421, p=0.658), but female rats did have more nose poke bouts than males in FR5-2 (Fig. 3F, Sex (F_(1,38)_=2.808, p=0.102), Test (F_(2,76)_=9.898, p<0.001) and Sex*Test interaction effects (F_(2,76)_=2.263, p=0.111), Post-hoc: males vs females: FR5-2: p=0.013). In addition, the number of nose poke bouts decreases the more FR5 tests both male and female rats perform (Females: FR5-3 vs FR5-1: p=0.037, FR5-3 vs FR5-2: p=0.025, Males: FR5-1 vs FR5-2: p=0.001, FR5-1 vs FR5-2: p=0.018), without affecting the mean duration of these bouts (Fig. 3G, Sex (F_(1,38)_=2.753, p=0.105), Test (F_(2,76)_=1.937, p=0.151) and Sex*Test interaction effects (F_(2,76)_=0.047, p=0.954) or the time-outs between them (Fig. 3H, Sex (F_(1,38)_=1.450, p=0.236), Test (F_(2,76)_=0.919, p=0.403) and Sex*Test interaction effects (F_(2,76)_=0.821, p=0.444)).

When comparing S1 and S2 of the rats that obtained two ejaculations in a test, we found no differences between ejaculatory series or sex on the number of nose pokes (Fig. 2I, Sex (F_(1,26.993)_=2.835, p=0.104), Test (F_(2,131.568)_=0.717, p=0.490), Series (F_(1,110.008)_=1.080, p=0.301) and Sex*Test*Series interaction effects (F_(7,116.327)_=0.683, p=0.686)) or number of crossings (Fig. 2K, Sex (F_(1,24.877)_=3.575, p=0.070), Test (F_(2,128.033)_=0.917, p=0.402), Series (F_(1,106.488)_=2.698, p=0.103) and Sex*Test*Series interaction effects (F_(7,112.698)_=0.907, p=0.504)). In contrast, we did find that male rats used longer latencies to poke in S2 in all FR5 tests compared to females (Fig. 2J, Sex (F_(1,35.569)_=20.094, p<0.001), Test (F_(2,136.477)_=0.212, p=0.809), Series (F_(1,119.501)_=190.899, p<0.001) and Sex*Test*Series interaction effects (F_(7,124.614)_=5.350, p<0.001), Post-hoc: Males vs females: S2: FR5-1: p=0.002, FR5-2: p<0.001, FR5-3: p<0.00). In addition, both male and female rats had longer poke latencies in S2 compared to S1 (Males: FR5-1: p<0.001, FR5-2: p<0.001, FR5-3: p<0.001; Females: FR5-1: p<0.001, FR5-2: p=0.002, FR5-3: p=0.009). Interestingly, this did not affect the latency to cross to the reward chamber in the same way. While no differences were found on the latency to cross between S1 and S2 in both male and females rats, females rats only had longer latencies to cross compared to males in S2 of FR5-2 (Fig. 2L, Sex (F_(1,34.128)_=10.292, p=0.003), Test (F_(2,142.523)_=2.451, p=0.090), Series (F_(1,122.099)_=0.217, p=0.642) and Sex*Test*Series interaction effects (F_(7,128.624)_=0.278, p=0.962), Post-hoc: Males vs females: S2: FR5-2: p=0.019).

#### 3.2.3 PR

Finally, the rats underwent three additional trials in the MCC test using a progressive ratio (PR) schedule. The first two PR tests (PR-1 and PR-2) were concluded either after 10 minutes of inactivity (no nose pokes) or when 30 minutes had passed since the start of the session. The final PR test (PR-3) was terminated under the same inactivity criterion of 10 minutes, but with an extended session duration of up to 60 minutes. While no differences in number of nose pokes were found in the FR1 and FR5 schedules, we now found that female rats poked generally more than male rats in the PR tests (Fig. 4A, Sex (F_(1,38)_=23.126, p<0.001), Test (F_(2,76)_=21.658, p<0.001) and Sex*Test interaction effects (F_(2,76)_=9.275, p<0.001), Post-hoc: Male vs Female, PR-1: p=0.016, PR-2: p=0.037, PR-3: p<0.001). In addition, the extra 30 minutes in the test box during PR-3 did not change the total number of nose pokes in males, but did significantly increase the number of pokes in females compared the PR-1 and PR-2 (Females: PR-3 vs PR-1: p<0.001, PR-3 vs PR-2: p<0.001). A similar effect was found on the latency to poke, with females returning faster to poking than males in all PR tests (Fig. 4B, Sex (F_(1,38)_=23.799, p<0.001), Test (F_(2,76)_=3.398, p=0.039) and Sex*Test interaction effects (F_(2,76)_=0.054, p=0.948), Post-hoc: Male vs Female, PR-1: p=0.001, PR-2: p<0.001, PR-3: p<0.001). The differences in nose poke behavior between male and female rats, however, did not result in differences in number of crossings (Fig. 4C, Sex (F_(1,38)_=4.604, p=0.038), Test (F_(2,76)_=1.642, p=0.200) and Sex*Test interaction effects (F_(2,76)_=0.859, p=0.428). Male and female rats crossed the same number of times towards the reward chamber in PR-1 and PR-2, and only in PR-3 females crossed more than males (Males vs females: PR-3: p=0.017). No differences in number of crossings were found between PR tests, nor did we find any differences (between sex or tests) on the latency to cross (Fig. 4D, Sex (F_(1,20.136)_=1.134, p=0.300), Test (F_(2,50.323)_=0.533, p=0.590) and Sex*Test interaction effects (F_(2,50.323)_=2.834, p=0.068).

**Figure 4.**
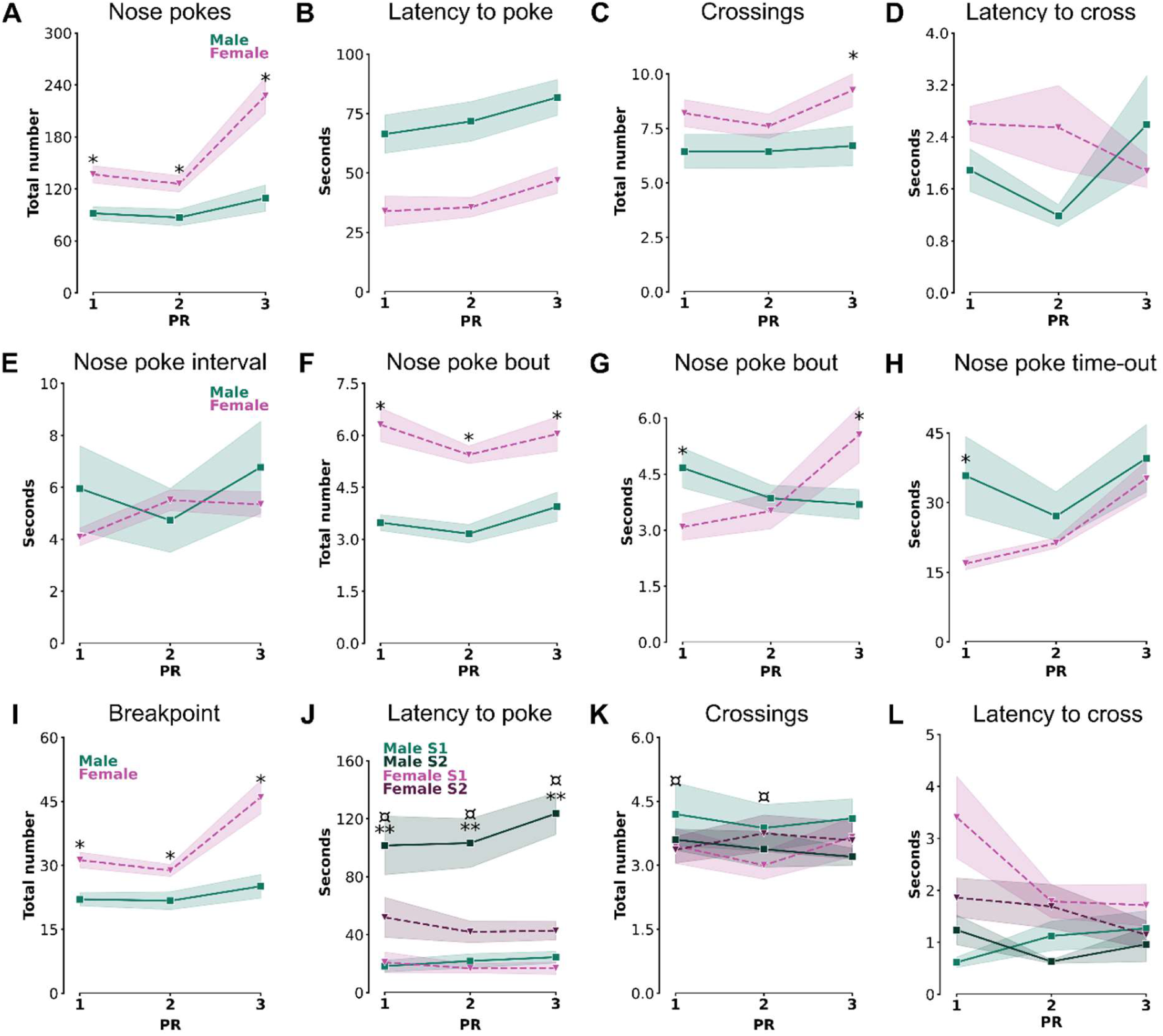
A) Total number of nose pokes, B) Latency to nose poke, C) Total number of crossings, and D) Latency to cross, E) Nose poke interval, F) Total number of nose poke bouts, G) Mean duration of nose poke bouts, H) Mean duration of nose poke time-outs, and I) Breakpoint during the complete test under a PR schedule, and J) Total number of nose pokes, K) Total number of crossings, and L) Latency to cross during the 1st (S1) and 2nd (S2) ejaculatory series under a PR schedule. Data are shown of male (green line) and female (purple dotted line) rats in A-D, and male (green line) and female (purple dotted line) rats per ejaculatory Series 1 (light color line) and Series 2 (dark color dotted line) in E-H. Shadows represent the SEM. * p≤ 0.05 difference between male and female rats in total test (A-D) or Series 1 (EH), ** p≤ 0.05 difference between male and female rats in Series 2, ^ p≤ 0.05 difference between Series 1 and 2 of female rats, ¤ p≤ 0.05 difference between Series 1 and 2 of male rats.

While again no sex or test differences were found in nose poke intervals (Fig. 4E, Sex (F_(1,38)_=0.370, p=0.547), Test (F_(2,76)_=1.213, p=0.303) and Sex*Test interaction effects (F_(2,76)_=1.867, p=0.162), female rats actually did poke with more nose poke bouts than male rats in all 3 PR tests (Fig. 4F, Sex (F_(1,38)_=39.754, p<0.001), Test (F_(2,76)_=2.713, p=0.073) and Sex*Test interaction effects (F_(2,76)_=20.698, p=0.501), Post-hoc: Females vs males: PR-1: p<0.001, PR-2: p<0.001, PR-3: p<0.001). While the mean duration of these nose poke bouts increased over the course of multiple PR tests in females, it decreased in males (Fig. 4G, Sex (F_(1,38)_=0.002, p=0.962), Test (F_(2,76)_=1.890, p=0.158) and Sex*Test interaction effects (F_(2,76)_=5.886, p=0.004), Post-hoc: Females vs males: PR-1: p=0.027, PR-3: p=0.009, Females: PR-3 vs PR-1: p=0.003, PR-3 vs PR-2: p=0.018). Interestingly, besides having longer nose poke bouts, male rats also took longer time-outs between these bouts than females in PR-1. This effect disappear across PR tests, when females increased their mean duration of time-outs while males remained stable across tests (Fig. 4H, Sex (F_(1,38)_=2.632, p=0.113), Test (F_(2,76)_=6.474, p=0.003) and Sex*Test interaction effects (F_(2,76)_=2.075, p=0.133), Post-hoc: Females vs males: PR-1: p=0.014, Females: PR-3 vs PR-1: p=0.005, PR-3 vs PR-2: p=0.044).

The benefit of a PR regime compared to FR is the use of the breakpoint parameter. The breakpoint is usually measured as the number of nose pokes required to open the door, after which the animal stops poking for a certain amount of time (in our study 10 minutes). To prevent the influences of sexual exhaustion in the follow-up tests, we were forced to end the test after 30 minutes (in PR1 and PR2) or 60 minutes (in PR3) if the natural breakpoint was not yet reached. The breakpoint is therefore, unfortunately, not directly comparable to breakpoints in other kinds of experiments. Still, it represents an interesting parameter, and we found that female rats had significantly higher breakpoints than males in all PR tests (Fig. 4D, Sex (F_(1,38)_=22.465, p<0.001), Test (F_(2,76)_=17.196, p<0.001) and Sex*Test interaction effects (F_(2,76)_=7.616, p<0.001). Males vs Females: PR-1: p=0.009, PR-2: p=0.041, PR-3: p<0.001). Interestingly, only in females, the extra 30 minutes in PR-3 resulted in significantly higher breakpoints compared to PR1 (p<0.001) and PR2 (p<0.001).

When comparing S1 with S2 in the PR test, it is expected in this kind of test that both males and female rats poked more frequently during S2 than S1, as it is a requirement of the PR test (Sex (F_(1,32.397)_=0.000, p=0.993), Test (F_(2,101.233)_=0.177, p=0.838), Series (F_(1,80.415)_=67.998, p<0.001) and Sex*Test*Series interaction effects (F_(7,87.551)_=0.856, p=0.545), Post-hoc: S1 vs S2: Females: PR-1: p<0.001, PR-2: p<0.001, PR-3: p<0.001, Males: PR-1: p=0.020, PR-2: p=0.002, PR-3: p=0.012). However, in addition to these increased number of nose pokes, we also found that both male and females rats took longer to return to nose poking after contact with the sexual stimulus in S2 than S1 (Fig. 4J, Sex (F_(1,28.077)_=19.704, p<0.001), Test (F_(2,97.415)_=0.507, p=0.604), Series (F_(1,74.913)_=171.058, p<0.001) and Sex*Test*Series interaction effects (F_(7,81.850)_=8.135, p<0.001), Post-hoc: S1 vs S2: Females: PR-1:p=0.002, PR-2: p=0.008, PR-3: p=0-006, Males: PR-1: p<0.001, PR-2: p<0.001, PR-3: p<0.001). In S2, males had longer latencies than females as well (PR-1: p<0.001, PR-2: p<0.001, PR-3: p<0.001). However, despite the increased number of nose pokes or longer latencies, they did not cross differently in S2 vs S1 (Fig. 4K, Sex (F_(1,27.792)_=0.980, p=0.331), Test (F_(2,101.900)_=0.149, p=0.861), Series (F_(1,82.136)_=0.849, p=0.360) and Sex*Test*Series interaction effects (F_(7,89.328)_=0.787, p=0.600)), nor did they have differences in latency to cross (Fig. 4L, Sex (F_(1,26.504)_=21.102, p<0.001), Test (F_(2,102.068)_=1.264, p=0.287), Series (F_(1,84.907)_=2.303, p=0.133) and Sex*Test*Series interaction effects (F_(7,91.154)_=1.593, p=0.147)). In addition, no differences in crossing behavior were found between male and female rats.

#### 3.2.4 Copulation behavior during all MCC tests

Upon successful nose poke series, the rats could sexually interact with their partner. Although male and female sexual behaviors are not directly comparable, it is still interesting to study how many behaviors each sex performs or receives in a MCC test. As shown in Table 2, we found that females received more ejaculations on average than males achieved in almost all FR1 test (FR1 tests: Sex: F_(1,38)_= 26.664, p<0.01, Test: F_(5,190)_=3.474, p=0.005 or Sex*Test interaction effects F_(5, 190)_=2.189, p=0.057, post-hoc: Male vs Female: FR1-1: p<0.001, FR1-2: p<0.001, FR1-3: p=0.013, FR1-4: p<0.001, FR1-5:p=0.022) and the 1st PR test (PR tests: Sex: F_(1,38)_= 6.336, p=0.016, Test: F_(2,76)_=0.622, p=0.539 or Sex*Test interaction effects F_(2,76)_=0.622, p=0.539, post-hoc: Male vs Female: PR-1: p=0.010), but not FR5 tests (FR5 tests: Sex: F_(1,38)_= 4.344, p=0.044, Test: F_(2,76)_=1.248, p=0.293 or Sex*Test interaction effects F_(2,76)_=0.153, p=0.858). While male rats needed time to gain sexual experience (FR1-2 vs FR1-5: p=0.004, FR1-2 vs FR1-6: p=0.004), females remained constant over the course of tests (they were paired with sexually experienced male rats). However, when looking at the total number of (received or performed) copulations, the males only show more copulations during the FR1-2 and less in the FR5-2 test compared to females, while during the PR tests, no differences were found (FR1 tests: Sex: F_(1,38)_= 0.237, p=0.629, Test: F_(5,190)_=1.219, p=0.302 or Sex*Test interaction effects F_(5, 190)_=2.577, p=0.028, post-hoc: Male vs Female: FR1-2: p=0.034; FR5 tests: Sex: F_(1,38)_= 5.688, p=0.022, Test: F_(2,76)_=0.152, p=0.859 or Sex*Test interaction effects F_(2,76)_=0.152, p=0.859, post-hoc: Male vs Female: FR5-2: p=0.043; PR tests: Sex: F_(1,38)_= 1.263, p=0.268, Test: F_(2,76)_=2.481, p=0.090 or Sex*Test interaction effects F_(2,76)_=0.294, p=0.746).

**Table 2.** Sexual behavioral outcomes during the motivation to continue copulation tests.

| Parameter | Sex | FR1 |  |  |  |  |  | FR5 |  |  | PR |  |  |
| --- | --- | --- | --- | --- | --- | --- | --- | --- | --- | --- | --- | --- | --- |
|  |  | 1 | 2 | 3 | 4 | 5 | 6 | 1 | 2 | 3 | 1 | 2 | 3 |
| Number of ejaculations | Female | 1.55±<br>0.15 | 1.8±<br>0.12 | 1.75±<br>0.12 | 1.75±<br>0.14 | 1.85±<br>0.11 | 1.75±<br>0.12 | 1.85±<br>0.08 | 1.6±<br>0.17 | 1.75±<br>0.1 | 1.85±<br>0.25 | 1.75±<br>0.25 | 1.85±<br>0.28 |
|  | Male | 0.7±<br>0.19* | 0.55±<br>0.18* | 1.15±<br>0.22* | 0.8±<br>0.2* | 1.3±<br>0.19* | 1.3±<br>0.21 | 1.45±<br>0.17 | 1.3±<br>0.19 | 1.5±<br>0.18 | 0.95±<br>0.2* | 1.15±<br>0.18* | 1.35±<br>0.26 |
| Number of copulations | Female | 12.8±<br>1.17 | 14.85±<br>1.66 | 14.35±<br>1.42 | 10.95±<br>1.53 | 11.9±<br>1.02 | 11.85±<br>1.01 | 10.8±<br>1.07 | 9.9±<br>1.26 | 11.45±<br>1.28 | 12.1±<br>1.25 | 12.25±<br>1.1 | 15.25±<br>1.97 |
|  | Male | 8.45±<br>2.22 | 8.9±<br>2.71* | 13.45±<br>2.69 | 12.05±<br>2.47 | 13.5±<br>1.98 | 14.7±<br>2.51 | 15.7±<br>2.48 | 15.25±<br>2.57* | 15.2±<br>1.83 | 10.65±<br>1.59 | 10.75±<br>1.53 | 12.2±<br>1.9 |
| Latency to 1<sup>st</sup> copulatory behavior | Female | 189.25±<br>57.82 | 115.36±<br>51.66 | 68.75±<br>17.85 | 219.42±<br>115.48 | 38.13±<br>9.69 | 53.33±<br>27.65 | 52.4±<br>15.61 | 322.02±<br>143.21 | 37.85±<br>6.66 | 159.7±<br>101.18 | 44.05±<br>15.44 | 242.77±<br>127.53 |
|  | Male | 806.74±<br>171.45* | 979.87±<br>177.96* | 773.57±<br>188.09* | 740.87±<br>165.39* | 416.05±<br>147.16* | 539.47±<br>169.69* | 365.55±<br>143.33 | 478.91±<br>154.4 | 329.23±<br>133.16 | 490.11±<br>173.71 | 383.87±<br>162.47 | 499.21±<br>174.69 |
| Ejaculation latency | Female | 591.28±<br>101.72 | 409.91±<br>110.94 | 365.26±<br>96.73 | 349.23±<br>128.19 | 303.44±<br>90.35 | 261.0±<br>84.26 | 224.94±<br>37.83 | 484.34±<br>133.55 | 294.78±<br>69.36 | 379.24±<br>115.19 | 422.47±<br>134.87 | 500.6±<br>133.49 |
|  | Male | 1260.69±<br>143.16* | 1449.63±<br>122.31* | 935.58±<br>164.76* | 1116.08±<br>160.49* | 660.19±<br>153.81 | 762.6±<br>161.44* | 689.69±<br>134.97* | 727.31±<br>148.83 | 616.83±<br>122.59 | 965.39±<br>160.0* | 767.19±<br>149.65 | 923.39±<br>177.74* |
| Number of paracopulatory behaviors | Female | 111.9±<br>12.84 | 74.0±<br>10.52 | 71.15±<br>12.87 | 55.7±<br>8.74 | 56.85±<br>9.46 | 58.7±<br>8.52 | 30.25±<br>2.59 | 33.45±<br>3.55 | 33.9±<br>6.13 | 46.55±<br>4.18 | 47.65±<br>5.61 | 39.95±<br>5.69 |
| Number of paracopulatory behaviors during the nose poke phase with closed doors | Female | 65.15±<br>7.72 | 23.8±<br>4.48 | 15.4±<br>3.46 | 16.4±<br>3.31 | 12.75±<br>2.4 | 13.3±<br>2.6 | 7.25±<br>2.06 | 4.35±<br>1.69 | 2.6±<br>0.8 | 10.0±<br>2.89 | 11.95±<br>2.4 | 7.6±<br>1.36 |
| Lordosis quotient | Female | 105.07±<br>2.23 | 101.61±<br>2.03 | 104.17±<br>1.81 | 101.3±<br>2.66 | 100.71±<br>1.73 | 102.05±<br>2.77 | 102.94±<br>2.47 | 104.37±<br>2.61 | 100.57±<br>1.27 | 100.84±<br>1.15 | 102.99±<br>2.31 | 105.39±<br>3.54 |
| Lordosis quotient upon received copulations | Female | 99.01±<br>0.75 | 98.15±<br>1.19 | 98.9±<br>0.73 | 97.51±<br>2.2 | 97.76±<br>1.82 | 97.1±<br>2.17 | 98.05±<br>0.97 | 98.27±<br>0.94 | 98.44±<br>0.71 | 99.1±<br>0.63 | 98.58±<br>0.95 | 98.49±<br>1.23 |
| Lordosis score | Female | 2.72±<br>0.07 | 2.77±<br>0.04 | 2.75±<br>0.06 | 2.74±<br>0.07 | 2.8±<br>0.06 | 2.8±<br>0.06 | 2.64±<br>0.04 | 2.66±<br>0.07 | 2.64±<br>0.06 | 2.82±<br>0.05 | 2.77±<br>0.06 | 2.63±<br>0.1 |

Note: \* $p \leq 0.05$ difference between male and female rats in the test.

Despite the lack of meaningful differences in the number of (received or performed) copulations, the latency to the 1^st^ copulatory behavior was significantly higher in all FR1 tests in males than females, but not in the FR5 or PR tests (FR1 tests: Sex: F_(1,38.114)_= 28.548, p<0.001, Test: F_(5,189.231)_=2.529, p=0.030 or Sex*Test interaction effects F_(5, 189.231)_=1.232, p=0.296, post-hoc: Male vs Female: FR1-1: p<0.001, FR1-2: p<0.001; FR1-3: p<0.001, FR1-4: p=0.004, FR1-5: p=0.041, FR1-6: p=0.008; FR5 tests: Sex: F_(1,38)_= 3.540, p=0.068, Test: F_(2,76)_=3.973, p=0.023 or Sex*Test interaction effects F_(2,76)_=0.506, p=0.605; PR tests: Sex: F_(1,38)_= 4.471, p=0.041, Test: F_(2,76)_=1.046, p=0.356 or Sex*Test interaction effects F_(2,76)_=0.084, p=0.920). Also when calculating the latency to the 1^st^ ejaculation from the 1^st^ copulatory behavior (mount or intromission), the males remained slower than the females in most of FR1 tests, PR test and FR5-1 (FR1 tests: Sex: F_(1,38.078)_= 27.783, p<0.001, Test: F_(5,189.169)_=6.911, p<0.001 or Sex*Test interaction effects F_(5, 189.169)_=2.493, p=0.033, post-hoc: Male vs Female: FR1-1: p<0.001, FR1-2: p<0.001, FR1-3: p=0.002, FR1-4: p<0.001, FR1-6: p=0.007; FR5 tests: Sex: F_(1,38)_= 8.029, p=0.007, Test: F_(2,76)_=1.676, p=0.194 or Sex*Test interaction effects F_(2,76)_=0.713, p=0.493, post-hoc: Male vs Female: FR5-1: p=0.005; PR tests: Sex: F_(1,38)_= 9.206, p=0.004, Test: F_(2,76)_=0.454, p=0.637 or Sex*Test interaction effects F_(2,76)_=0.486, p=0.617, post-hoc: Male vs Female: PR-1: p=0.006, PR-3: p=0.044). Only males showed a significant longer ejaculation latency in FR1-1 vs FR1-5 (p=0.001) and FR1-6 (p=0.014).

With regard to female sexual behavior, we found that the number of paracopulatory behaviors declined over the course of testing (FR1 tests: Test: F_(5,95)_= 7.684, p<0.001, post-hoc: FR1-1 vs FR1-2: p=0.011, FR1-1 vs FR1-3: p=0.005, FR1-1 vs FR1-4: p<0001, FR1-1 vs FR1-5: p<0.001, FR1-1 vs FR1-6: p<0.001; FR5 tests: Test: F_(2,38)_= 0.235, p=0.792; PR tests: Test: F_(2,38)_=0.806, p=0.454), which was mainly a result of performing less paracopulatory behavior in the period in which they can nose poke to gain access to the male (FR1 tests: Test: F_(5,95)_= 30.466, p<0.001, post-hoc: FR1-1 vs FR1-2: p<0.001, FR1-1 vs FR1-3: p<0.001, FR1-1 vs FR1-4: p<0001, FR1-1 vs FR1-5: p<0.001, FR1-1 vs FR1-6: p<0.001; FR5 tests: Test: F_(2,38)_= 3.379, p=0.045, post-hoc: FR5-1 vs FR5-3: p=0.042; PR tests: Test: F_(2,38)_=0.899, p=0.415). However, in respect to the lordosis behavior, we found that the females showed a constant and high level of receptivity from the 1^st^ to the last MCC test. No differences were found in lordosis quotient (independent of whether or not it was calculated upon all lordosis responses or only those related to received copulatory behaviors) (FR1 tests: Test: F_(5,95)_= 0.746, p=0.591; FR5 tests: Test: F_(2,35.979)_= 0.045, p=0.986; PR tests: Test: F_(2,37.956)_=0.742, p=0.483). The lordosis score was also always very high and consistent (FR1 tests: Test: F_(5,94.054)_= 1.160, p=0.335; FR5 tests: Test: F_(2,36.050)_= 0.964, p=0.391; PR tests: Test: F_(2,36.603)_=3.146, p=0.055).

### 3.3 Motivation comparison between tests

As previously mentioned, sexual behaviors in males and females differ substantially, making direct comparisons challenging. Therefore, we did not perform statistical analyses to examine sex differences in sexual behaviors during the copulation test. However, all behavioral outcomes are presented in Table S1. The primary objective of the copulation tests was to explore potential correlations between specific parameters and the outcomes of the MCC test. To acclimate the rats to this new testing paradigm, the copulation test was conducted three times. Male rats were tested in a single-compartment box, while females were tested in a two-compartment box. Comparative analyses were performed using data from the third copulation (COP3) test only.

Regarding comparisons with the sexual incentive motivation (SIM) test, we correlated SIM1 with the first three FR1 tests, SIM2 with the last three FR1 tests and FR5 tests, and SIM3 with the PR tests. A correlation was deemed meaningful only if the same effect was consistently observed across several tests, and p-values were below 0.01.

The results showed that none of the parameters of the MCC test were correlated with either the SIM or copulation test in male rats, nor did any parameter of SIM1 or SIM3 correlate with a parameter of the copulation test. Only the time spent with the control incentive in SIM2 correlated positively with the latency to 1^st^ copulatory behavior in the copulation test (*ρ*=0.588, p=0.006).

Similar to males, we found no correlations between parameters of the MCC and SIM tests in female rats. In addition, we also found no correlation between parameters of the FR1 tests and the copulation test. However, we found some random correlations appearing between some parameters. Although we do not interpret them as particularly relevant, we still found that the inter poke interval in all FR5 tests negatively correlated with the number of intromissions in the copulation test (FR5-1: *ρ*=-0.495, p=0.027, FR5-2: *ρ*=-0.623, p=0.003, FR5-3, *ρ*=-0.696, p<0.001), something that was also true for (only) the PR-3 test (*ρ*=-0.699, p<0.001). Furthermore, we found that the Latency to poke in FR5 tests were negatively correlated with the total number of ejaculations received in the copulation test (FR5-1: *ρ*=-0.587, p=0.006, FR5-2: *ρ*=-0.587, p=0.006, FR5-3: *ρ*=-0.469, p=0.037). In the PR test, we found a negative correlation between the total number of copulatory behaviors in PR tests and the mean duration of time-out in the copulation test (PR-2: *ρ*=-0.568, 0.009, PR-3, *ρ*=-0.568, p=0.009).

Moreover, we did find some correlations between the SIM and copulation test outcomes in female rats. The preference score of SIM1 positively correlated with the total number of intromissions (*ρ*=0.559, p=0.010) and the total number of sexual bouts (*ρ*=0.609, p=0.004). The time spent with the control incentive, on the other hand, did negatively correlate with the mean duration of the time-outs (*ρ*=0.579, p=0.007) and total number of copulations (*ρ*=-0.620, p=0.004). In SIM3, a negative correlation was found between the preference score and the contact-return latency after intromissions (*ρ*=-0.782, p=0.004) and a positive correlation between the time spent with the control incentive and the contact-return latency after intromissions (*ρ*=0.845, p=0.001). No other correlations have been detected.

## 4. Discussion

In this study, we aimed to establish a paradigm that categorizes sexual motivation into two distinct components: sexual incentive motivation and the motivation to continue copulation. The Sexual Incentive Motivation (SIM) test was designed to effectively measure incentive-driven motivation, while the Motivation to Continue Copulation (MCC) test, based on nose-poking behavior, was developed to assess the drive to sustain copulatory activity. The validity of the MCC test is underscored by findings in male rats, where the latency to initiate nose poking upon returning to the nose-poke compartment increased during the second ejaculatory series compared to the first. This effect was particularly pronounced under higher-effort nose-poke schedules, such as fixed ratio (FR) 5 and progressive ratio (PR), though it was already evident under the lower-effort FR1 schedule. A similar pattern was observed in female rats tested under an FR5 schedule. Notably, the increased latency to nose poke was not accompanied by reductions in the total number of nose pokes or crossings to access the sexual partner. Furthermore, our findings reveal that sexually experienced rats exhibit greater motivation to continue copulation compared to sexually naïve rats, and that baseline nose-poking behaviors differ between male and female rats.

Our findings suggest that latency to nose poke may be a more sensitive and reliable parameter for assessing the motivation to continue copulation compared to the total number of nose pokes or crossings. While the number of nose pokes and crossings reflects the overall effort exerted to gain access to a sexual partner, these measures did not consistently show significant changes across ejaculatory series or between different effort schedules. In contrast, latency to nose poke demonstrated clear and reproducible differences, particularly between the first and second ejaculatory series, even under varying levels of effort (e.g., FR1, FR5, and PR schedules). This suggests that latency captures a distinct aspect of motivation, potentially reflecting the initial drive or motivation to reinitiate sexual behavior after ejaculation. Additionally, latency is less likely to be influenced by external factors, such as the physical effort required to perform repeated nose pokes or crossings, making it a more direct measure of motivational state.

The longer latencies to nose pokes observed in both male (FR1, FR5, and PR) and female (FR5) rats during the second ejaculatory series suggest a nuanced shift in the motivation to continue copulation after ejaculation. From an evolutionary perspective, this decline in motivation is logical, particularly for males, as reproductive success diminishes with subsequent ejaculations due to a reduction in available sperm cells (Austin and Dewsbury, 1986; Ågmo, 1976). Interestingly, our findings also show that males required fewer copulations to achieve a second ejaculation, aligning with previous studies reporting fewer intromissions and shorter ejaculation latencies during the second ejaculatory series (Bermant, 1964; Chu and Ågmo, 2015; Dewsbury, 1969; Larsson, 1959; Price, 1980; Sachs, 1976). While this might initially seem contradictory to the observed decline in motivation, it is likely explained by physiological factors, such as increased penile sensitivity, which facilitate quicker ejaculation with fewer copulations. Moreover, the prolonged post-ejaculatory intervals that typically follow multiple ejaculations (Dewsbury, 1969; Larsson, 1959; Rodríguez-Manzo and Fernández-Guasti, 1994; Sachs, 1976) appear to play a significant role in the reduced motivation observed in the MCC test. Notably, the most pronounced effects were evident in the FR5 and PR schedules, which require greater effort to access the sexual stimulus and are therefore more sensitive measures of the motivation to continue copulation. However, even in the low-effort FR1 schedule, a decline in crossings during the second ejaculatory series was observed in the later tests, suggesting that this decrease in motivation is a robust phenomenon, regardless of the effort required.

Since sexual interaction is highly rewarding, one might contrarily expect the motivation to continue copulation to increase rather than diminish in series 2. Nevertheless, this pattern might reflect the economic concept of diminished marginal utility, where the value of a reward decreases with subsequent exposure. In this context, the first ejaculation might likely be more rewarding in that it satisfies an immediate motivational drive, temporarily reducing the perceived value of the drive to continue copulating for a second ejaculation. This reduction in perceived value may override the expectation that continuing mating interaction would lead to greater motivation, as the initial reinforcement provided by the first ejaculation appears sufficient to induce a reward state. Curiously, while several studies have reported that ejaculations induced a conditioned place preference (Tenk et al., 2009; Ågmo and Berenfeld, 1990), no study has yet shown whether the rewarding nature of ejaculations is diminished in the second ejaculatory series relative to the first. This remains an open question for further research.

Female rats, in contrast, did not consistently exhibit significant differences in their motivation to continue copulation across ejaculatory series. While increased latencies to nose poke were observed in FR5 tests, the visually apparent increases in PR tests did not reach statistical significance, and no differences were detected in FR1 tests. Unlike males, females cannot control the type of stimulation they receive during copulation; however, previous research has shown that their behavioral responses, such as contact-return latencies, increase with the intensity of stimulation (Brandling-Bennett et al., 1999; Erskine, 1985; Erskine, 1989; Krieger et al., 1976; Meerts et al., 2014; Zipse et al., 2000). Similarly, in seminatural environments, inter-lordosis intervals have been found to be longer following ejaculations compared to mounts (Chu and Ågmo, 2014). Despite these variations in behavioral responses, they do not appear to have significant biological consequences for reproductive success. Fertility is unaffected by the number of ejaculations received (Chu and Ågmo, 2014) and instead, the ability to pace sexual interactions plays a critical role in creating the physiological conditions necessary for successful fertilization (Coopersmith and Erskine, 1994; Erskine, 1989). Furthermore, it has been demonstrated that control over the timing of stimulation, rather than the stimulation itself, is what induces a conditioned place preference (Arzate et al., 2011). Taken together, these findings, along with the fact that females have much shorter post-ejaculatory intervals (especially now we changed males after an ejaculation), may explain why their motivation to continue copulation remains consistently high across ejaculatory series under low-effort conditions. However, this motivation may appear to waver when the effort required to gain access increases, as seen in FR5 and PR schedules.

This study investigated differences between sexually naïve and experienced rats. During the MCC test training phase, rats successfully learned the task using cheese rewards and continued to perform well when the reward was switched to a sexual stimulus, validating the approach for sexually naïve subjects. No significant differences were observed in the number of nose pokes or crossings between naïve and experienced rats, regardless of sex. However, sexually naïve male rats exhibited longer latencies to nose poke during the first two FR1 tests, resulting in fewer copulations and ejaculations despite identical opportunities for sexual interaction. These findings align with previous research showing that sexual experience alters male copulatory patterns, leading to faster initiation of copulation, higher intromission ratios, and increased ejaculation numbers (Dewsbury, 1969; Larsson, 1959). Sexual experience also influences the rewarding aspects of sexual interactions, as naïve males develop a conditioned place preference for intromissions alone, whereas experienced males require ejaculation to perceive the same reward (Tenk et al., 2009). This suggests that sexual interaction involves a learning process, with experience stabilizing the motivation to continue copulation. Consistent with this, sexually naïve males exhibited immediate sexual incentive motivation, which strengthened with experience, as reflected in increased preference scores in SIM3 compared to SIM1 (Lopez et al., 1999; Ågmo, 2003).

In female rats, no effects of sexual experience on sexual incentive motivation were observed. However, sexually naïve females showed longer latencies during initial FR1 tests in the 1^st^ ejaculatory series, which decreased over time, suggesting an increase in motivation to continue copulation with repeated exposure. Previous studies have reported no significant differences in sexual behavior between naïve and experienced females (Oyem et al., 2025), though the impact of sexual training on female sexual performance remains debated (Blaustein et al., 2009; Meerts et al., 2016; Meerts et al., 2014; Meerts et al., 2015; Oyem et al., 2025). These findings suggest that sexual experience plays a critical role in shaping motivation and behavior in male rats, while its effects on females may be more subtle and context-dependent.

This study also examined potential sex differences in sexual motivation. Both males and females demonstrated sexual incentive motivation in sexually naïve and experienced states. Females showed higher preference scores than males in the naïve condition, while males exhibited increased incentive motivation after gaining sexual experience. Sex differences were also observed in nose poke behavior. Males had higher latencies to poke during the first two FR1 tests and in FR5 overall, while females performed more nose pokes, achieved higher breakpoints, and engaged in more nose poke bouts in PR. During the first ejaculatory series, males performed more nose pokes and crossings in FR1, but in the second series, males exhibited longer latencies to poke than females in FR5. At first glance, these findings might suggest that females are more motivated to continue copulation than males, while sexually experienced males exhibit higher levels of incentive motivation. However, it is more likely that males and females have inherently different baselines in the MCC test, complicating direct comparisons. As shown in Table S1, females engage in significantly more sexual bouts due to their increased paracopulatory behaviors compared to the mounts and intromissions performed by males. Additionally, females receive more copulations despite having longer inter-intromission intervals and lower intromission ratios. These differences, while intriguing, do not necessarily reflect disparities in mating efficiency between the sexes. To draw meaningful conclusions, it is essential to evaluate parameters within each sex group rather than directly comparing males and females. Although the MCC test was initially expected to facilitate sex comparisons, the results suggest otherwise. For example, females were unaffected by the males’ post-ejaculatory intervals during the test, giving them more time to engage in nose poking and likely explains their higher levels of nose poke behavior and shorter latencies. However, these results should be interpreted cautiously, as they may also reflect inherent differences in the sexual behavior and baseline activity of each sex rather than differences in motivation alone.

The final aim of this study was to evaluate whether specific parameters from the standard copulation (COP) test could serve as measures of sexual incentive motivation or the motivation to continue copulation or the other way around. Our results did not indicate any relevant and consistent correlation between parameters of tests. The lack of correlations between the SIM and MCC tests supports the hypothesis that sexual motivation can be divided into distinct components, with the SIM test specializing in measuring sexual incentive motivation and the MCC test focusing on the motivation to continue copulation. In our previous studies we suggested that parameters such as sexual bouts and time-outs might reflect motivation to continue copulation (Huijgens et al., 2021a; Oyem et al., 2025), but the current findings do not support this hypothesis. Instead, these parameters appear to reflect temporal patterns of copulatory behavior that are independent of motivational states measured in a conditioning box. Still, it is important to realize that the MCC test insists on a fixed time limit for copulation during each access trial (15 seconds), which may influence the observed behaviors. For example, the timing of separation at the end of a trial could coincide with different behavioral states. In males, separation might occur during a natural time-out or in the middle of a mount bout, potentially affecting the latency to nose poke in the subsequent trial. This variability in behavioral state at the time of separation could explain the lack of correlation between nose poke behavior in the MCC test and sexual bouts or time-outs in the COP test. To address this issue in future studies, it may be beneficial to allow copulation in the MCC test to continue until the end of a mount bout before returning the rat to the nose poke compartment. Despite being practically difficult, this adjustment could provide a clearer understanding of the relationship between copulatory patterns and motivational states.

While this study provides valuable insights into the distinct components of sexual motivation and their measurement, several limitations should be acknowledged to contextualize the findings and guide future research. One limitation lies in the inability to measure a meaningful breakpoint in the progressive ratio (PR) test due to the nature of rat copulatory behavior. Male rats can continue copulating after multiple ejaculations, with short post-ejaculatory intervals of approximately five minutes (Beach, 1966; Chu and Ågmo, 2015, 2014; Hegstad et al., 2020), making it impractical to determine a natural breakpoint before sexual exhaustion. Sexual exhaustion, typically reached after 7–8 ejaculations (Fernandez-Guasti and Rodriguez-Manzo, 2003; Hull and Dominguez, 2007; Karen and Barfield, 1975; Mas et al., 1995; Ventura-Aquino and Fernandez-Guasti, 2013), likely represents the true breakpoint. The Coolidge effect, where male rats resume copulation with a new partner even after exhaustion, further suggests that sexual exhaustion and the reinstatement of sexual activity are closely tied to the motivation to continue copulation (Rodriguez-Manzo, 1999; Ventura-Aquino et al., 2016; Ventura-Aquino et al., 2018). However, due to the prolonged recovery period required after sexual exhaustion, the repeated testing design of this study did not allow for PR tests until exhaustion. Instead, testing was limited to 30–60 minutes per session. Future studies should explore the effects of mating until exhaustion in the MCC test to provide deeper insights into sexual motivation and behavior.

Another limitation involves the hormone-dependent nature of female rat sexual behavior, which follows a 4–5-day estrous cycle. Daily testing would require hormone injections, potentially leading to a build-up of hormone levels and unnatural conditions. To address this, females were hormonally primed and tested every fourth day, allowing hormone levels to return to baseline. While this approach provided a more natural setting, it limited the ability to conduct daily testing, which is typically beneficial for establishing stable nose poke behavior in conditioning paradigms. Individual fluctuations in nose poke behavior were observed in both females and males, the latter of which were tested every other day to prevent sexual exhaustion. Interestingly, the highest activity was observed on the fourth test in males, consistent with previous findings that male rats exhibit cyclicity in activity, with peak behavior occurring every four days (Shulman and Spritzer, 2014). This suggests that both male and female rats may have inherent cycles of activity that could influence test outcomes.

Additionally, a key limitation of this study is the lack of experimental manipulations to directly alter motivation in the MCC test. Testing the effects of pharmacological, hormonal, or environmental manipulations on motivation would provide stronger evidence for the specificity and sensitivity of the MCC test in measuring the motivation to continue copulation. Disruptions in general and sexual motivation are often observed in individuals with depression, PTSD, and other psychiatric conditions, where both incentive and consummatory aspects of motivation may be affected. The SIM and MCC paradigms offer a valuable tool set for disentangling these components, providing a clearer understanding of how specific motivational deficits contribute to psychiatric dysfunctions (Association, 2013; Everitt et al., 2008; Stone et al., 2008). Moreover, these paradigms could be adapted to study the effects of pharmacological, hormonal, or environmental manipulations on sexual motivation, but also other motivation-driven concepts, offering insights into potential therapeutic strategies for restoring (sexual) motivation in clinical populations.

## 5. Conclusion

In conclusion, this study provides novel insights into the distinct components of sexual motivation and their measurement in male and female rats, with significant implications for both basic and translational research. By employing the SIM and MCC tests, sexual motivation can effectively be divided into sexual incentive motivation and the motivation to continue copulation, respectively. The findings highlight the influence of sexual experience on male and female sexual behavior. Despite the absence of manipulations to directly alter motivation in the MCC test, the study establishes a robust framework for investigating sexual motivation to continue copulation and provides a foundation for future studies to explore the behavioral, physiological, and neurobiological mechanisms underlying sexual motivation in both sexes. Importantly, the translational relevance of these paradigms extends beyond basic research, as they can inform studies on psychiatric conditions where disruptions in incentive and consummatory motivation are common.

## Author contributions

JCO: Experimental design, data gathering, behavioral annotation, data curation, analysis, writing – original draft.

PTH: Experimental design, behavioral annotation, writing – review and editing

JM: behavioral annotation

RH: Experimental design, methodology, supervision, writing – review and editing, funding acquisition.

EMSS: Experimental design, methodology, data curation, Programming/software, analysis, supervision, writing – original draft, funding acquisition.

## Acknowledgments

Financial support was received from Helse Nord (HNF1443-19) and the AKM fund of UiT The Arctic University of Norway. We sincerely appreciate Amalie Hofmeyer, Carina Sørensen, Remi Osnes, and Hallvard Haugen for the excellent care of the experimental animals. We especially thank Lorenzo Ragazzi and Izabela Szkotak for injecting our rats with progesterone in the morning. We also extend our gratitude to Truls Traasdahl, Thomas Nermo and the local workshop for their skillful design and construction of our behavioral boxes. Finally, we would also like to thank Ida Johannessen for her assistance with the behavioral annotations of the male copulation test.

## Declaration of interests

The authors declare no competing interests.

## Notes

### Competing Interest Statement

The authors have declared no competing interest.

## References

Arzate DM, Portillo W, Rodriguez C, Corona R, Paredes RG. Extended paced mating tests induces conditioned place preference without affecting sexual arousal. Horm. Behav., 2011; 59: 674–80.

Association AP. Diagnostic and Statistical Manual of Mental Disorders, Fifth Edition (DSM-5). American Psychiatric Association, 2013: 947.

Austin D, Dewsbury DA. Reproductive capacity of male laboratory rats. Physiol. Behav., 1986; 37: 627–32.

Beach FA. Sexual attractivity, proceptivity, and receptivity in female mammals. Horm. Behav., 1976; 7: 105–38.

Beach FA. Sexual behavior in the male rat. Science, 1966; 153: 769–70.

Bermant G. Effects of Single and Multiple Enforced Intercopulatory Intervals on the Sexual Behavior of Male Rats. J. Comp. Physiol. Psychol., 1964; 57: 398–403.

Bermant G. Response Latencies of Female Rats during Sexual Intercourse. Science, 1961; 133: 1771–3.

Bialy M, Bogacki-Rychlik W, Przybylski J, Zera T. The Sexual Motivation of Male Rats as a Tool in Animal Models of Human Health Disorders. Front. Behav. Neurosci., 2019; 13.

Blaustein JD, Farrell S, Ghavami G, Laroche J, Mohan G. Non-intromissive mating stimuli are sufficient to enhance sexual behaviors in ovariectomized female rats. Horm. Behav., 2009; 55: 404–11.

Brandling-Bennett EM, Blasberg ME, Clark AS. Paced mating behavior in female rats in response to different hormone priming regimens. Horm. Behav., 1999; 35: 144–54.

Chu X, Ågmo A. Sociosexual Behaviors of Male Rats (Rattus norvegicus) in a Seminatural Environment. J. Comp. Psychol., 2015.

Chu X, Ågmo A. Sociosexual behaviours in cycling, intact female rats (Rattus norvegicus) housed in a seminatural environment. Behaviour, 2014; 151: 1143–84.

Coopersmith C, Erskine MS. Influence of paced mating and number of intromissions on fertility in the laboratory rat. J. Reprod. Fertil., 1994; 102: 451–8.

Cummings JA, Becker JB. Quantitative assessment of female sexual motivation in the rat: Hormonal control of motivation. J. Neurosci. Methods, 2012; 204: 227–33.

Dewsbury DA. Copulatory behaviour of rats (Rattus norvegicus) as a function of prior copulatory experience. Anim. Behav., 1969; 17: 217–23.

Dewsbury DA. A Quantitative Description of Behavior of Rats during Copulation. Behaviour, 1967; 29: 154–C.

Ellingsen E, Ågmo A. Sexual-incentive motivation and paced sexual behavior in female rats after treatment with drugs modifying dopaminergic neurotransmission. Pharmacol. Biochem. Behav., 2004; 77: 431–45.

Erskine MS. Effects of paced coital stimulation on estrus duration in intact cycling rats and ovariectomized and ovariectomized-adrenalectomized hormone-primed rats. Behavioral neuroscience, 1985; 99: 151.

Erskine MS. Solicitation behavior in the estrous female rat: a review. Horm. Behav., 1989; 23: 473–502.

Everitt BJ. Sexual motivation: a neural and behavioural analysis of the mechanisms underlying appetitive and copulatory responses of male rats. Neurosci. Biobehav. Rev., 1990; 14: 217–32.

Everitt BJ, Belin D, Economidou D, Pelloux Y, Dalley JW, Robbins TW. Review. Neural mechanisms underlying the vulnerability to develop compulsive drug-seeking habits and addiction. Philos. Trans. R. Soc. Lond. B Biol. Sci., 2008; 363: 3125–35.

Everitt BJ, Stacey P. Studies of instrumental behavior with sexual reinforcement in male rats (Rattus norvegicus): II. Effects of preoptic area lesions, castration, and testosterone. J. Comp. Psychol., 1 87; 101: 407–19.

Fernandez-Guasti A, Rodriguez-Manzo G. Pharmacological and physiological aspects of sexual exhaustion in male rats. Scand. J. Psychol., 2003; 44: 257–63.

French D, Fitzpatrick D, Law OT. Operant investigation of mating preference in female rats. J. Comp. Physiol. Psychol., 1972; 81: 226–32.

Gelez H, Greggain-Mohr J, Pfaus JG, Allers KA, Giuliano F. Flibanserin treatment increases appetitive sexual motivation in the female rat. J. Sex. Med., 2013; 10: 1231–9.

Hardy DF, Debold JF. Effects of mounts without intromission upon the behavior of female rats during the onset of estrogen-induced heat. Physiol. Behav., 1971; 7: 643–5.

Hegstad J, Huijgens PT, Houwing DJ, Olivier JDA, Heijkoop R, Snoeren EMS. Female rat sexual behavior is unaffected by perinatal fluoxetine exposure. Psychoneuroendocrinology, 2020; 120: 104796.

Heijkoop R, Huijgens PT, Snoeren EMS. Assessment of sexual behavior in rats: The potentials and pitfalls. Behavioural Brain Research, 2018a; 352: 70–80.

Heijkoop R, Huijgens PT, Snoeren EMS. Assessment of sexual behavior in rats: The potentials and pitfalls. Behav. Brain Res., 2018b; 352: 70–80.

Hodos W. Progressive ratio as a measure of reward strength. Science, 1961; 134: 943–4.

Huijgens PT, Guarraci FA, Olivier JDA, Snoeren EMS. Male rat sexual behavior: Insights from inter-copulatory intervals. Behav. Processes, 2021a; 190: 104458.

Huijgens PT, Heijkoop R, Snoeren EMS. Sexual Incentive Motivation. In Paredes RG, Portillo W, Bedos M, editors. Animal Models of Reproductive Behavior. Springer US: New York, NY, 2023: 197–210.

Huijgens PT, Heijkoop R, Snoeren EMS. Silencing and stimulating the medial amygdala impairs ejaculation but not sexual incentive motivation in male rats. Behav. Brain Res., 2021b; 405: 113206.

Huijgens PT, Heijkoop R, Vanderschuren L, Lesscher HMB, Snoeren EMS. CaMKIIa+ neurons in the bed nucleus of the stria terminalis modulate pace of natural reward seeking depending on internal state. Psychopharmacology (Berl.), 2024.

Hull EM, Bazzett TJ, Warner RK, Eaton RC, Thompson JT. Dopamine receptors in the ventral tegmental area modulate male sexual behavior in rats. Brain Res., 1990; 512: 1–6.

Hull EM, Dominguez JM. Sexual behavior in male rodents. Horm. Behav., 2007; 52: 45–55

Karen LM, Barfield RJ. Differential rates of exhaustion and recovery of several parameters of male rat sexual behavior. J. Comp. Physiol. Psychol., 1975; 88: 693–703.

Kippin TE, Sotiropoulos V, Badih J, Pfaus JG. Opposing roles of the nucleus accumbens and anterior lateral hypothalamic area in the control of sexual behaviour in the male rat. Eur. J. Neurosci., 2004; 19: 698–704.

Krieger MS, Orr D, Perper T. Temporal patterning of sexual behavior in the female rat. Behavioral Biology, 1976; 18: 379–86.

Larsson K. Experience and maturation in the development of sexual behavior in male puberty rat. Behaviour, 1959; 14: 101–7.

Le Moene O, Agmo A. Modeling Human Sexual Motivation in Rodents: Some Caveats. Front. Behav. Neurosci., 2019; 13: 187.

Lopez HH, Olster DH, Ettenberg A. Sexual motivation in the male rat: the role of primary incentives and copulatory experience. Horm Behav, 1999; 36: 176–85.

Mas M, Fumero B, Fernandez-Vera JR, Gonzalez-Mora JL. Neurochemical correlates of sexual exhaustion and recovery as assessed by in vivo microdialysis. Brain Res., 1995; 675: 13–9.

Matthews TJ, Grigore M, Tang L, Doat M, Kow LM, Pfaff DW. Sexual reinforcement in the female rat. J. Exp. Anal. Behav., 1997; 68: 399–410.

Meerts SH, Park JH, Sekhawat R. Sexual experience modulates partner preference and mPOA nitric oxide synthase in female rats. Behav Neurosci, 2016; 130: 490–9.

Meerts SH, Schairer RS, Farry-Thorn ME, Johnson EG, Strnad HK. Previous sexual experience alters the display of paced mating behavior in female rats. Horm. Behav., 2014; 65: 497–504.

Meerts SH, Strnad HK, Schairer RS. Paced mating behavior is affected by clitoral-vaginocervical lidocaine application in combination with sexual experience. Physiology & Behavior, 2015; 140: 222–9.

Oyem JC, Heijkoop R, Snoeren EM. The temporal copulatory patterns of female rat sexual behavior. Behav. Processes, 2025: 105148.

Pfaus JG, Mendelson SD, Phillips AG. A correlational and factor analysis of anticipatory and consummatory measures of sexual behavior in the male rat. Psychoneuroendocrinology, 1990; 15: 329–40.

Portillo W, Paredes RG. Sexual incentive motivation, olfactory preference, and activation of the vomeronasal projection pathway by sexually relevant cues in non-copulating and naive male rats. Horm. Behav., 2004; 46: 330–40.

Price EO. Sexual-Behavior and Reproductive Competition in Male Wild and Domestic Norway Rats. Anim. Behav., 1980; 28: 657–67.

Richardson NR, Roberts DC. Progressive ratio schedules in drug self-administration studies in rats: a method to evaluate reinforcing efficacy. J. Neurosci. Methods, 1996; 66: 1–11.

Rodriguez-Manzo G. Blockade of the establishment of the sexual inhibition resulting from sexual exhaustion by the Coolidge effect. Behav. Brain Res., 1999; 100: 245–54.

Rodríguez-Manzo G, Fernández-Guasti A. Reversal of sexual exhaustion by serotonergic and noradrenergic agents. Behavioural Brain Research, 1994; 62: 127–34.

Sachs BD. Functional Analysis of Masculine Copulatory Behavior in the Rat. Advances in the Study of Behavior, 1976; 7: 91–154.

Sewalem J, Kassaw C, Anbesaw T. Sexual dysfunction among people with mental illness attending follow-up treatment at a tertiary hospital, Jimma University Medical Center: A cross-sectional study. Front Psychiatry, 2022; 13: 999922.

Shulman LM, Spritzer MD. Changes in the sexual behavior and testosterone levels of male rats in response to daily interactions with estrus females. Physiol. Behav., 2014; 133: 8–13.

Snoeren EM, Bovens A, Refsgaard LK, Westphal KG, Waldinger MD, Olivier B, Oosting RS. Combination of testosterone and vardenafil increases female sexual functioning in sub-primed rats. J. Sex. Med., 2011a; 8: 989–1001.

Snoeren EM, Chan JS, de Jong TR, Waldinger MD, Olivier B, Oosting RS. A new female rat animal model for hypoactive sexual desire disorder; behavioral and pharmacological evidence. J. Sex. Med., 2011b; 8: 44–56.

Snoeren EM, Lehtimaki J, Ågmo A. Effect of dexmedetomidine on ejaculatory behavior and sexual motivation in intact male rats. Pharmacol. Biochem. Behav., 2012; 103: 345–52.

Snoeren EM, Ågmo A. The incentive value of males’ 50-kHz ultrasonic vocalizations for female rats (*Rattus norvegicus*). J. Comp. Psychol., 2014; 128: 40–55.

Snoeren EMS. Female Reproductive Behavior. Curr. Top. Behav. Neurosci., 2019; 43: 1–44.

Stone EA, Lin Y, Quartermain D. A final common pathway for depression? Progress toward a general conceptual framework. Neurosci. Biobehav. Rev., 2008; 32: 508–24.

Tenk CM, Wilson H, Zhang Q, Pitchers KK, Coolen LM. Sexual reward in male rats: effects of sexual experience on conditioned place preferences associated with ejaculation and intromissions. Horm. Behav., 2009; 55: 93–7.

Vega Matuszcyk J, Larsson K, Eriksson E. The selective serotonin reuptake inhibitor fluoxetine reduces sexual motivation in male rats. Pharmacol. Biochem. Behav., 1998; 60: 527–32.

Ventura-Aquino E, Banos-Araujo J, Fernandez-Guasti A, Paredes RG. An unknown male increases sexual incentive motivation and partner preference: Further evidence for the Coolidge effect in female rats. Physiol. Behav., 2016; 158: 54–9.

Ventura-Aquino E, Fernandez-Guasti A. Reduced proceptivity and sex-motivated behaviors in the female rat after repeated copulation in paced and non-paced mating: effect of changing the male. Physiol. Behav., 2013; 120: 70–6.

Ventura-Aquino E, Fernandez-Guasti A, Paredes RG. Hormones and the Coolidge effect. Mol. Cell. Endocrinol., 2018; 467: 42–8.

Ventura-Aquino E, Paredes RG. Animal Models in Sexual Medicine: The Need and Importance of Studying Sexual Motivation. Sex Med Rev, 2017; 5: 5–19.

Waldinger MD. Psychiatric disorders and sexual dysfunction. Handb. Clin. Neurol., 2015; 130: 469–89.

Zipse LR, Brandling-Bennett EM, Clark AS. Paced mating behavior in the naturally cycling and the hormone-treated female rat. Physiol. Behav., 2000; 70: 205–9.

Ågmo A. Male rat sexual behavior. Brain Res. Brain Res. Protoc., 1997; 1: 203–9.

Ågmo A. The number of spermatozoa in spontaneous ejaculates of rats. J. Reprod. Fertil., 1976; 48: 405–7.

Ågmo A. On the concept of sexual arousal: a simpler alternative. Horm. Behav., 2008; 53: 312–4; author reply 9–22.

Ågmo A. Unconditioned sexual incentive motivation in the male Norway rat (Rattus norvegicus). J. Comp. Psychol., 2003; 117: 3–14.

Ågmo A, Berenfeld R. Reinforcing properties of ejaculation in the male rat: role of opioids and dopamine. Behav. Neurosci., 1990; 104: 177–82.

Ågmo A, Laan E. The Sexual Incentive Motivation Model and Its Clinical Applications. J. Sex Res., 2023; 60: 969–88.

Ågmo A, Laan E. Sexual incentive motivation, sexual behavior, and general arousal: Do rats and humans tell the same story? Neurosci. Biobehav. Rev., 2022; 135: 104595.

